# Salt Inducible Kinases as Novel Notch Interactors in the Developing *Drosophila* Retina

**DOI:** 10.1101/786004

**Authors:** H. Bahar Şahin, Sercan Sayın, Kuyaş Buğra, Arzu Çelik

## Abstract

Developmental processes require strict regulation of proliferation, differentiation and patterning for the generation of final organ size. Aberrations in these fundamental events are critically important in understanding tumorigenesis and cancer progression. Salt inducible kinases (Siks) are evolutionarily conserved genes involved in diverse biological processes, including salt sensing, metabolism, muscle and skeletal development. Recent findings implicate SIKs in tumor suppression or progression. However, their role in development remains largely unknown.

Using a sensitized tumor model in the *Drosophila* eye, we show that perturbations of Sik function exacerbates tumor-like tissue overgrowth and metastasis. Furthermore, we show that both *Drosophila Sik* genes, *Sik2* and *Sik3*, are required for proper eye development. We propose that an important target of Siks may be the Notch pathway, as we demonstrate epistasis between Siks and Notch pathway members and identify putative phosphorylation motifs on Notch, Delta and Fringe. Finally, we investigate Sik expression in the developing retina and show that Sik2 is expressed in all photoreceptors in close proximity to cell junctions, while Sik3 appears to be expressed specifically in R3/R4 cells in the developing eye. Combined, our data suggest that *Sik* genes are important in tissue specification, growth, and that their dysregulation may contribute to tumor formation.

## Introduction

The generation of tissues and the establishment of cellular diversity in multicellular organisms rely on cell-cell communication during development. One of the prominent and evolutionarily-conserved signaling pathways that regulate specification and patterning of tissues is the Notch signaling pathway. Notch has been established as a key regulator of development in both vertebrates and invertebrates (reviewed in (Zhang et al 2018b)). Interestingly, many of the signaling pathways, including the Notch pathway, are involved in cell fate determination and have been implicated in promoting or suppressing cancer when mutated or misregulated (Bossuyt et al 2009).

The *Drosophila* eye has been used as a model to study development, patterning, and tumor growth. The eye starts to develop in the second instar larval stage, from the eye-antennal imaginal disc, a single-layer epithelium. The posterior side of the disc is specified as eye primordium, marked by Notch expression, while the anterior is specified as antenna and is marked by EGFR expression (Kumar & Moses 2001) (**Supp. Figure 1A**). During the late second instar stage, a second Notch signal is critical to establish the dorsal-ventral (D-V) polarity of the eye disc, expressed by the organizer midline cells. The Notch receptor is induced by its ligand Delta on the dorsal side, and Serrate on the ventral side (Dominguez & de Celis 1998). The role of the glycosyltransferase Fringe is to modify Notch and inhibit its activation in the ventral half except the midline (Sato & Tomlinson 2007) (**Supp. Figure 1B**). In the late third instar stage, a differentiation wave known as the morphogenetic furrow (MF) moves from posterior to the anterior of the disc (**Supp. Figure 1C**) leaving differentiated cells behind. Photoreceptors (PRs) and accessory cells are specified sequentially from undifferentiated cells, with each step requiring Notch signaling (Voas & Rebay 2004). Once the neural and nonneural cells complete differentiation and planar cell polarity is established, the PRs start to elongate in the apical-basal direction and change their morphology, while being firmly attached to each other via AJs that contains Cadherin protein (reviewed in (Tepass & Harris 2007)). This coordinated differentiation and organization of cells gives rise to the formation of an adult eye that is composed of ~800 camera-like ommatidia, each contains eight PRs, supporting cone, pigment, and bristle cells (reviewed in (Şahin & Çelik 2013)).

Salt-inducible kinases (Siks) represent an evolutionarily conserved family of proteins that belong to the AMP-activated protein kinase (AMPK) superfamily. The two fly homologs (Sik2 and Sik3) show the same protein organization and high similarity with their human counterparts (SIK1, SIK2, SIK3) (**Supp. Figure 2**). The kinase domains are well conserved (>85% similarity) and point mutation in this domain, homologous to the K170 residue of fly Sik2, lead to kinase-dead allele (Choi et al 2011).

Siks are Ser/Thr kinases and they also have a number of phosphorylation sites. Sik activity is regulated by two main signals; Lkb-1, a general tumor suppressor, and PKA, an effector of second messenger cAMP. Siks are not catalytically active until phosphorylated by Lkb-1 on their T-loop (Lizcano et al 2004). Phosphorylation by PKA is inhibitory, and mutation of the target serine to an alanine results in constitutively active Sik alleles, such as S1032A of fly Sik2 or S563A of fly Sik3 (**Supp. Figure 2, 4**) (Screaton et al 2004). The common downstream effectors of Siks are CREB-regulated transcription coactivator proteins (CRTCs / TORCs), HDACs and IRS-1; (Horike et al 2003).

Siks are involved in diverse biological processes, including sugar and lipid metabolism (Zhang et al 2018a), neuronal and glial survival (Sasaki et al 2011) (Kuser-Abali et al 2013), autophagy (Yang et al, 2013) and regulation of the circadian clock (Hayasaka et al 2017). Siks have been shown to regulate body size in *C. elegans* (Maduzia et al 2005), and were linked to establishment of cell polarity and regulation of tissue growth in *Drosophila* (Parsons et al 2017). Additionally, both *Drosophila* Sik2 and Sik3 were shown to interact with the Hippo pathway to regulate tissue growth and size (Wehr et al 2012) and cancer progression (reviewed in (Du et al 2016)). Siks are also implicated in diseases such as diabetes (Sall et al 2017), epilepsy (Proschel et al 2017) and in various cancer types (Ahmed et al 2010) (Bon et al 2015) (Liu et al 2016) (Zohrap et al 2018). Given their proposed roles in mitotic mechanisms and as nutrient-sensors, Siks have been suggested to be the link between diet and tumorigenesis and cancer progression (Hirabayashi & Cagan 2015) (Amara et al 2017). Although these kinases have been extensively studied in metabolism and homeostasis, Sik functions during development are less clear.

Here, we make use of a *Drosophila* tumor model in the retina to investigate whether Siks can induce tumor progression. Our results indicate that Siks are expressed in the developing eye disc, and Sik function is required for proper retinal fate determination and eye size. Furthermore, we show that Sik activity is epistatic with Notch activity, and that misregulation of Sik activity promotes Notch-associated tumor-like growth of the eye tissue. This, together with a proteome-wide Sik motif search, suggest that Notch pathway members are direct downstream targets of the Siks.

## Results

### Sik Proteins Regulate Growth in *Drosophila* Tumor Models

Our previous experiments in mammalian cell culture indicated that Sik proteins are involved in cancer progression (Zohrap et al 2018). Thus, we addressed whether they are able to do so in *in vivo* tumor models and employed a *Drosophila* Notch pathway-dependent ocular tumor model. Specifcally, overexpression of the Notch ligand Delta using *ey-GAL4* induces eye overgrowth and sensitizes the animal to tumorigenesis (**Figure 1A, Supp. Figure 1C**) (Dominguez & de Celis 1998) (Ferres-Marco et al 2006). “Eyeful” flies, which overexpress Delta and concurrently carry the *GS88A8* mutation (deregulation of the epigenetic silencers *lola* (*CG12052*) and *psq* (*CG2368*)), for example, display further eye enlargement and tumorigenesis (Ferres-Marco et al 2006) (Bossuyt et al 2009). Therefore, we tested Sik function both in the “sensitized” (*ey-GAL4, UAS-Delta/CyO*) and “eyeful” (*ey-GAL4, UAS-Delta, GS88A8/CyO*) backgrounds by crossing in the *ey-Gal4* driver that expresses the UAS target specifically in the embryonic eye primordia and in the third instar larval stage before the morphogenetic furrow (MF) (Kumar & Moses 2001) and the different Sik transgenes.

**Figure 1.**
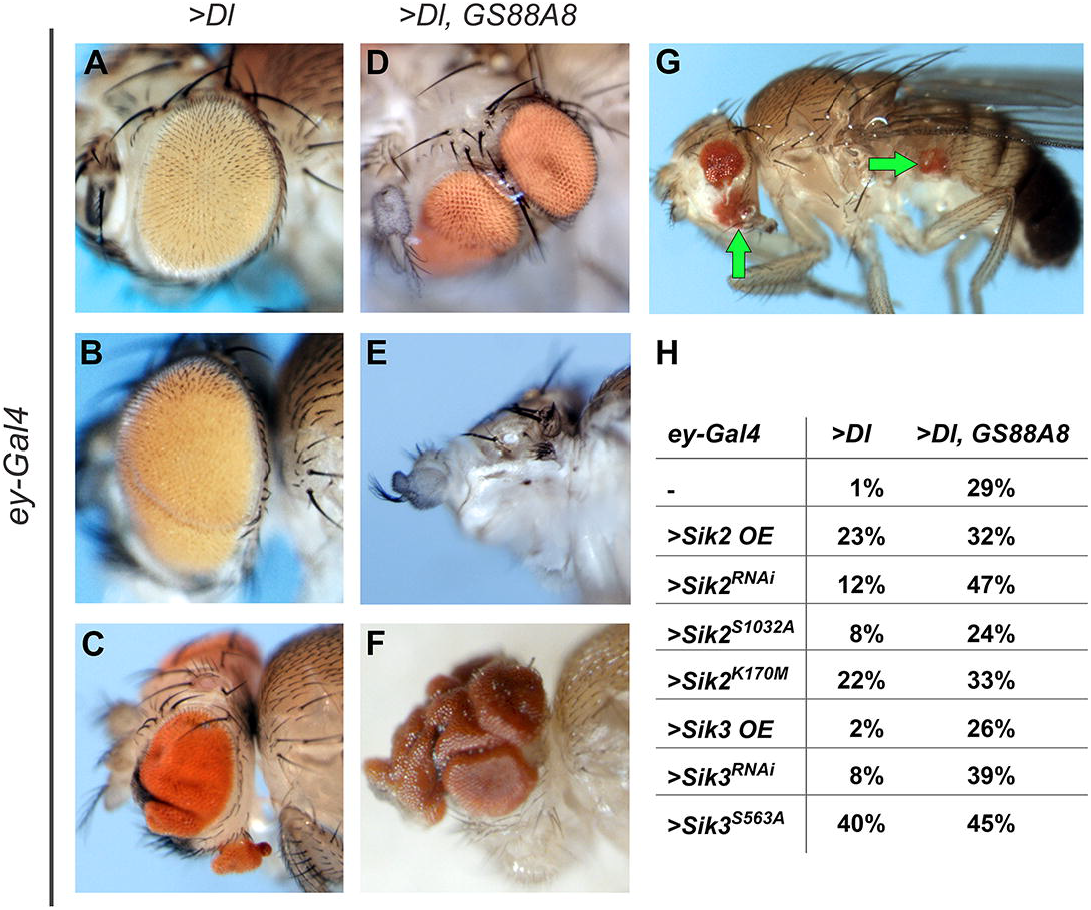
The tumor-like effect of salt inducible kinases (Siks). **A-G)** Examples of “sensitized” and “eyeful” crosses’ progenies. **A-C)** Examples of flies chosen from the sensitized background (*ey-Gal4 > Dl*) **A)** Fly from sensitized background showing slight eye overgrowth, accepted as normal / not affected. **B)** An example of eye overgrowth and folding, accepted as affected. **C)** An example of ectopic eye formation. **D-G)** Examples of flies chosen from the eyeful background (*ey-Gal4 > Dl*, *GS88A8*; *GS88A8* mutation disrupting epigenetic regulator *lola* and *pipsqueak* genes) **D)** An example of eye duplication; two independent eyes are formed on one side of the head. **E)** An example of eye-to-antenna differentiation. **F)** An example of excessive eye growth. **G)** A representative example of metastasis. The green arrows indicate proximal and distal metastasis of eye tissue. **H)** Sensitized (*ey-Gal4 > Dl*) and eyeful (*ey-Gal4>Dl, GS88A8*) flies were crossed with the Sik transgenes: wild type control, crosses with *W^1118^* (-), Sik2 overexpression (>*Sik2* OE), Sik2 knock-down (>*Sik2^RNAi^*), Sik2 constitutively active (>*Sik2^S1032A^*), Sik2 kinase-dead (>*Sik2^K170M^*), Sik3 overexpression (>*Sik3* OE), SIK3 knock-down (>*Sik3^RNAi^*), and Sik3 constitutively active (>*Sik3^S563A^*). For each cross, normal eyes (similar to the parent, like in **A**) and affected eyes (eye folding or overgrowth, duplicated eye or ectopic eye tissue, eye loss or eye-to-antenna differentiation, and metastasis, like in **B-G**) in the progeny were quantified. The ratio of cumulative number of affected eyes over the total eye number is presented in the table **H**. The detailed analysis is shown in **Supp. Table 1.** The full genotypes are listed in **Supp. Table 2**. Anterior is to the left, ventral is down.

All genetic combinations that were tested affected eye development to some extent. Affected eyes showed different phenotypes including tumor-like overgrowth, incidences of ectopic or duplicated eyes, total loss of eye tissue, and metastasis that originated from the dissemination of transformed cells from the developing eye to the body (**Figure 1A-G**). For each genetic combination, the ratio of affected eyes was quantified by analyzing and counting each eye independently. Eyes that resembled the parents were considered as not affected; any deviation from the parental phenotype was considered to be affected. In the sensitized background, Sik2 overexpression (*Sik2* OE) affected 23% of the eyes, Sik2 knock-down (*Sik2^RNAi^*) 12% of the eyes, Sik2 constitutively active (*Sik2^S1032A^*) 8% of the eyes, and Sik2 kinase-dead (*Sik2^K170M^*) affected 22% of the eyes. While Sik3 overexpression (*Sik3* OE) did not cause any eye phenotype different from the control, Sik3 knock-down (*Sik3^RNAi^*) affected 8% of eyes, (**Figure 1A-C,H**, **Supp. Table 1**). The ratio of affected eyes obtained in the eyeful background were in accordance with the ratio in sensitized background (**Figure 1D-F,H**, **Supp. Table 1**). The results show that, independent of increase or decrease, any modulation of Sik activity enhanced both tumor-like phenotypes and eye loss phenotypes, indicating blocked eye development. Unlike the above-mentioned combinations, constitutively active Sik3 (*Sik3^S563A^*) in the sensitized and eyeful backgrounds was mostly lethal. The survivors showed a higher percentage of phenotype, 40% and 45% of the eyes respectively; also, extreme cases of tissue defects, including differentiation problems of eye and antenna (**Figure 1E**) and rare instances of distal metastasis of eye tissue to the body (**Figure 1G**) were detected.

Thus, depletion or overexpression of Sik enhanced tumor-like growth in the eye. The ratio of affected eyes suggests that Sik2 plays a more critical role in this process than Sik3.Constitutively active Sik3 on the other hand, behaves more like wild type Sik2. Additionally, the observed eye-to-antenna transformation phenotypes suggest a possible role for Sik in retinal fate determination.

### Siks have a Role in the Developing Eye

To understand if Siks are required for normal patterning of the eye disc, Sik activity was specifically altered via two strong drivers. In this experiment, in addition to the *ey-Gal4* driver described in the previous experiment, we introduced *lGMR-Gal4* to drive expression in all PRs during larval stages and throughout adulthood (Moses & Rubin 1991), and *UAS-Dicer2* to increase the RNAi effect.

Modulating Sik protein levels with eye-specific drivers disrupted the eye structure both at the structural and cellular level. Overexpression of Sik2 (*Sik2* OE) and Sik3 (*Sik3* OE), as well as Sik3 knock-down (*Sik3^RNAi^*) caused defects in organization of ommatidia, lens and bristle cells and supernumerary bristles as shown by light microscopy and scanning electron microscopy at the structural level (**Figure 2A - B′′, E - F**′′). Sik2 knock-down (*Sik2^RNAi^*) not only affected the cellular organization, but also drastically decreased eye size with full penetrance (**Figure 2C - D**). Furthermore, it resulted in duplication of the antenna in some instances (**data not shown**). Duplicated antennae were also observed in eye imaginal discs (**Figure 2C**′′). Surprisingly, the Sik2 mutant (*Sik2^Δ41^*) did not exhibit lethality or any obvious defect in the eye (**data not shown**).

**Figure 2.**
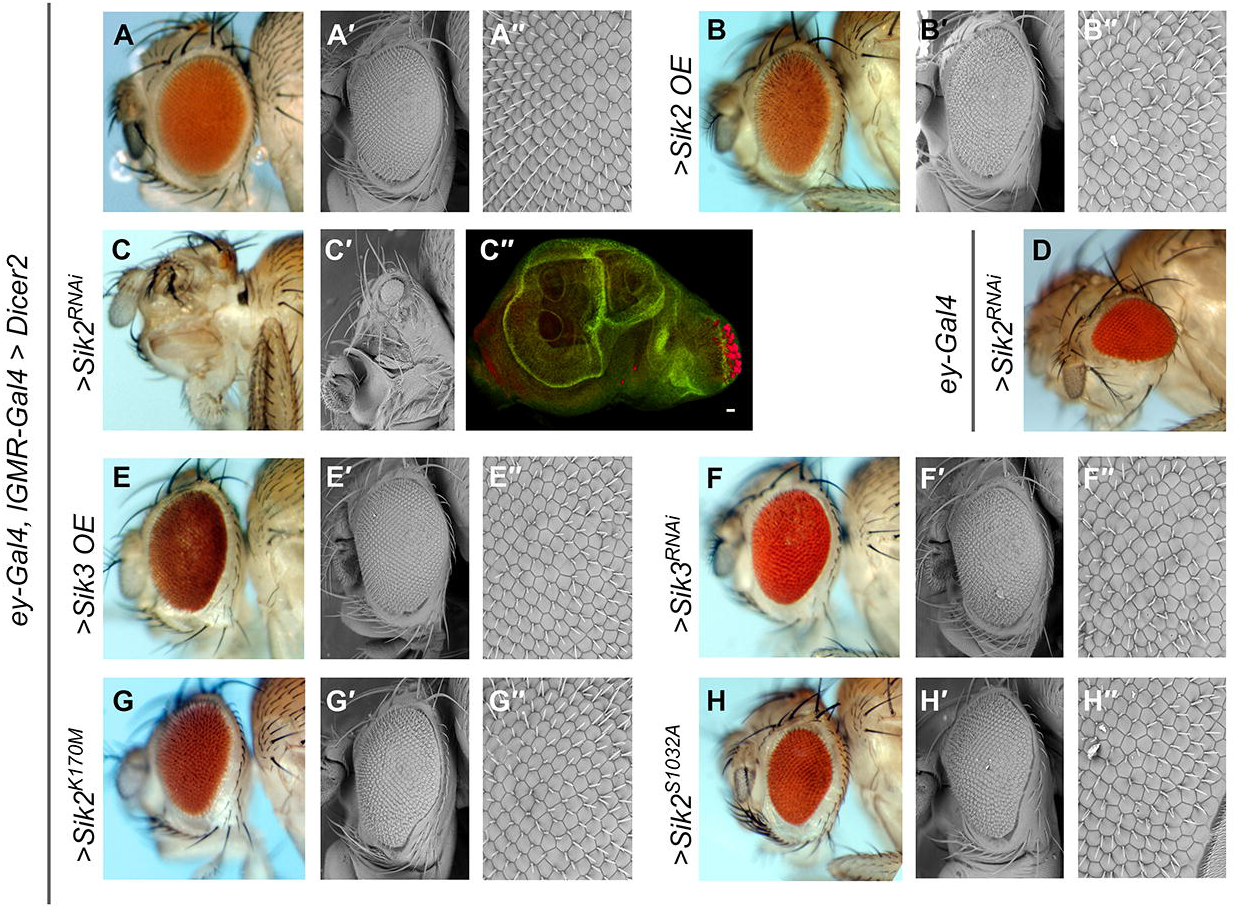
Eye morphology of Sik mutants. Images in **A**, **B**, **C**, **D**, **E, F, G, H** represent light microscopy, **A**′, **A**′′, **B**′, **B**′′, **C**′, **E**′, **E**′′**, F**′, **F**′′**, G**′, **G**′′**, H**′, **H**′′ scanning electron microscopy, **C**′′, confocal microscopy images. In all cases except **D**, the flies are in “*UAS-Dicer / +; ey-GAL4, lGMR-GAL4 / +*” background. **A**-**A**′′) Control (driver crossed with wild type, *W^1118^*). **B**-**B**′′) Sik2 overexpression (>*Sik2* OE). **C**-**C**′′) Sik2 knock-down (>*Sik2^RNAi^*) **C**′′) Third instar larval eye disc. Red is elav (neurogenic), green is Delta (non-specific stain, background signal). Scale bar 20 μm. **D**) Sik2 knock-down by *ey-*Gal4 driver and two copies of *UAS-Sik2^RNAi^*. **E**-**E**′′) SIK3 overexpression (>*Sik3* OE). **F**-**F**′′) Sik3 knock-down (>*Sik3^RNAi^*). **G-G**′′) Sik2 kinase-dead (>*SIK2^K170M^*). **H-H**′′) Sik2 constitutively active (>*Sik2^S1032A^*). The full genotypes are listed in **Supp. Table 2**. In all pictures, anterior is to the left, ventral is down.

Sik3 null mutants (*Sik3*^*Δ109*^) are lethal at an early stage, most probably due to a metabolic problem (Teesalu et al 2017). Thus, Sik3 null mutant eyes could only be investigated by generating eye-specific post-mitotic clones. We created Sik3 null mutant clones at the larval stage both in the entire eye, selecting against the apoptotic gene *hid* (**Supp. Figure 3A′′**), and in eye clones by employing the MARCM technique (**Supp. Figure 3B′**). As observed in the adult retinae, loss of Sik3 in clonal regions (white regions are *Sik3*^*Δ109*^ null mutant clones) did not visibly affect the eye structure in either experiment.

These results suggest that both Sik2 and Sik3 activity at a correct level are essential for proper eye development. While excess Sik2 or Sik3, or reduced Sik3, disrupts lens and bristle cell organization, reduction in Sik2 level prevents the development of the retina. The fact that null mutants have normal eye development suggests that either there is a compensation mechanism between Sik2 and Sik3 or RNAi transgenes knock-down both homologs. To overcome the compensation and off-target possibilities, point mutations were employed. Ectopic expression of a kinase-dead form of Sik2 (*Sik2^K170M^*) and a constitutively active form of Sik2 (*Sik2^S1032A^*) resulted in cone and bristle cell defects, similar to the phenotypes observed in other Sik overexpression and knock-down backgrounds. Additionally, these flies displayed a “notch” on the ventral side of the eye (**Figure 2G-H**′′**, data not shown**) that is not present in controls (**Figure 2A-A**′′). Furthermore, stronger knock-down of Sik2 (*Sik2^RNAi^*) with two copies of the *RNAi* allele via the *ey-Gal4* driver led to total loss of the ventral half of the eye (**Figure 2D**).

These ventral eye defects suggest a misregulation of dorsal-ventral (D-V) polarity during development. It is long known that the activation of Notch receptor in the eye disc midline cells by Delta and Serrate from dorsal and ventral sides respectively, establishes the D-V polarity and tissue development on the two sides at early larval stages (Dominguez & de Celis 1998) (**Supp. Figure 1B**). These observations suggest a possible interaction of Siks with the Notch signaling pathway. To strengthen this hypothesis, we investigated Sik function in the mechanosensory system, where macrochaetae are generated in a Notch-dependent manner. Ectopic expression of kinase-dead Sik2 (*Sik2^K170M^*), constitutively active Sik2 (*Sik2^S1032A^*), and constitutively active Sik3 (*Sik3^S563A^*) resulted in misshaped macrocheate (**Supp. Figure 4B,C,E**). Sik2 overexpression (*Sik2* OE) not only affected the macrocheate shape, but also microcheate number and organization (**Supp. Figure 4 D**).

Taken together, these data indicate that, disrupting the Sik balance in different systems results in phenotypes resembling Notch pathway defects.

### Siks Interact with the Notch Pathway

Analysis of Siks in the eyeful and sensitized backgrounds and during normal development indicate a possible interaction with the Notch pathway. To better understand this interaction, we used *ey-Gal4* and *lGMR-Gal4* drivers together to overexpress or deplete Notch pathway components in combination with *Sik* transgenes.

We first tested the Notch ligand Delta, which is both expressed in the early L3 stage to establish D-V polarity, and also expressed at late L3, just after the MF (**Supp. Figure 1B,C**). Different from the first experiment, another strong eye driver and Dicer was added to the background. Like the “sensitized” background (**Figure 1A**), overexpression of Delta resulted in larger eyes (**Figure 3A**), and this phenotype was enhanced by adding the constitutively active (*Sik2^S1032A^*) and kinase-dead (*Sik2^K170M^*) forms of Sik2 in most of the cases, with incidences of tissue folding and necrosis (**Figure 3B,C**). In some cases, a total loss of the eye was observed (**data not shown**). Comparing with the modulation of Sik activity in the same background, which has a minor effect on eye size that was limited to the ventral eye (**Figure 2G,H**), it clearly indicates a genetic interaction. While Sik3 knock-down (*Sik3^RNAi^*) did not affect the phenotype (**Figure 3D**), constitutively active Sik3 (*Sik3^S563A^*), which rather resembles Sik2 alleles in the “sensitized” and “eyeful” backgrounds, resulted in phenotypes similar to constitutively active Sik2 (*Sik2^S1032A^*), including larger eyes with necrotic lenses (**data not shown**) and total eye loss (**Figure 3E**). Sik2 knock-down (*Sik2^RNAi^*) in combination with Delta overexpression led to a phenotype very similar to constitutively active Sik2 (*Sik2^S1032A^*); larger eyes and necrotic lenses (**Figure 3F**). Keeping in mind that Sik2 knock-down itself (*Sik2^RNAi^*) in the same background prominently blocked eye growth with full penetrance (**Figure 2C**), as the Delta phenotype rescued the Sik2 phenotype, this experiment suggests that Delta might be working downstream of Sik2.

**Figure 3.**
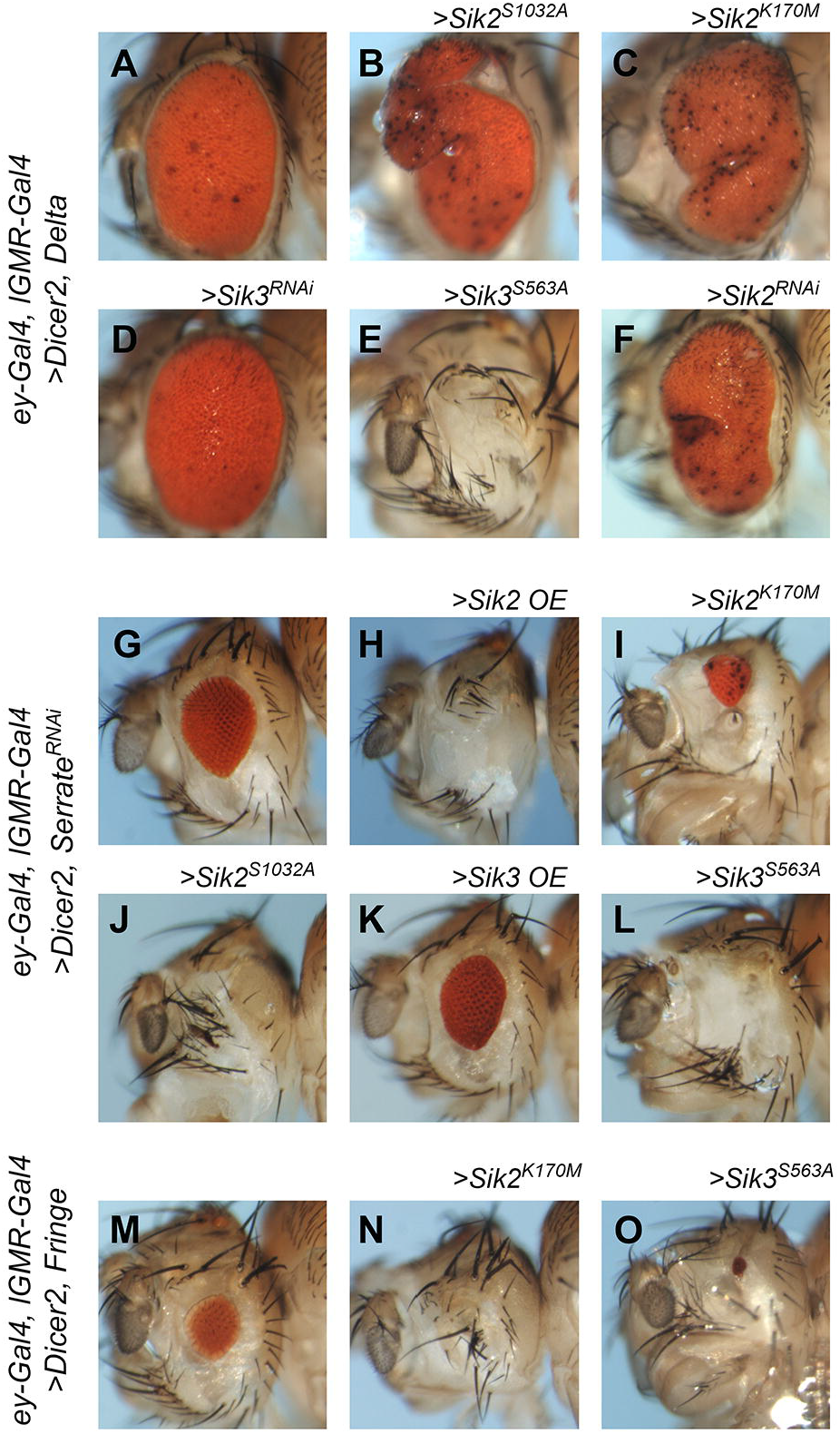
Genetic interaction between Sik tools and Notch pathway members. **A-F)** Delta overexpression with eye specific driver (*UAS-Dicer2 / UAS-Delta; ey-GAL4, lGMR-GAL4 / +*) in combination Siks. **A**) Control. **B**) Sik2 constitutively active (>*Sik2^S1032A^*). **C**) Sik2 kinase-dead (>*Sik2^K170M^*). **D**) Sik3 knock-down (>*Sik3^RNAi^*). **E**) Sik3 constitutively active (>*Sik3^S563A^*). **F**) Sik2 knock-down (>*Sik2^RNAi^*). **G-L)** Serrate knock-down with eye specific driver (*UAS-Dicer2 / +; ey-GAL4, lGMR-GAL4 / +; Serrate^RNAi^ / +*) in combination with Siks. **G**) Control **H**) Sik2 overexpression (>*Sik2* OE). **I**) Sik2 kinase-dead (>*Sik2^K170M^*). **J**) Sik2 constitutively active (>*Sik2^S1032A^*). **K**) Sik3 overexpression (>*Sik3* OE). **L**) SIK3 constitutively active (>*Sik3^S563A^*). **M-O)** Fringe overexpression by eye specific driver (*UAS-Dicer2 / +; ey-GAL4, lGMR-GAL4, UAS-Fringe / +*) in combination with Siks. **M**) Control. **N**) Sik2 kinase-dead (>*Sik2^K170M^*). **O**) Sik3 constitutively active (>*Sik3^S563A^*). All controls are crossed with wild type, *W^1118^*. The full genotypes are listed in **Supp. Table 2**. In all pictures, anterior is to the left, ventral is down.

Serrate is another Notch ligand, and it is expressed in the ventral side of the D-V border. Depletion of the Notch ligand Serrate using *Serrate^RNAi^* caused small eyes (**Figure 3G**). Sik2 overexpression (*Sik2* OE) in a Serrate-depleted background enhanced the small eye phenotype and even caused total loss of the eye in many instances (**Figure 3H**). Similar phenotypes were observed when Sik2 kinase-dead (*Sik2^K170M^*) or Sik2 constitutively active form (*Sik2^S1032A^*) were used (**Figure 3I,J**), suggesting a genetic interaction between Sik2 and Serrate. This epistatic interaction was confirmed using another *Serrate^RNAi^* allele (**data not shown**). Sik3 overexpression (*Sik3* OE) (**Figure 3K**) or depletion (**data not shown**) did not have any noticeable effect when combined with *Serrate^RNAi^*, just like in the Delta overexpression background (**Figure 3D**), whereas Sik3 constitutively active form (*Sik3^S563A^*) enhanced the small eye phenotype and resulted in total loss of the eye (**Figure 3L**). The constitutively active Sik3 phenotype resembled the phenotypes observed with Sik2 rather than Sik3 as in **Figure 1**, which might be attributed to a redundancy of Sik function as previously suggested (Wehr et al 2012).

The glycosyltransferase Fringe that modifies Notch to promote Notch-Delta interaction, is expressed in the ventral eye disc and suppresses Notch-Serrate interaction except for the dorsal-ventral border (**Supp. Figure 1B**). As previously reported *Fringe* overexpression results in smaller eyes, presumably by disrupting the balance of Notch activation during development (**Figure 3M**) (Dominguez et al 2004). The small eye phenotype of Fringe was greatly enhanced when combined with the kinase-dead form of Sik2 (*Sik2^K170M^*) or the constitutively active version of Sik3 (*Sik3^S563A^*) (**Figure 3N,O**), indicating a genetic interaction between Siks and Fringe. Sik3 overexpression on the other hand did not modify the Fringe overexpression phenotype at all (**data not shown**), like the Sik3 combinations above (**Figure 3D,K**).

Taken together, these observations suggest that there is epistasis between Notch pathway members and Siks, in particular Sik2. Since Sik is a protein kinase, we hypothesize that Sik2 targets and phosphorylates members of the Notch pathway directly, or indirectly through the effectors of the pathway.

In order to investigate if Notch pathway members could be directly targeted by Siks, we set out to find Sik phosphorylation motifs on Notch pathway members. First, we aimed to determine the amino acid sequence targeted by Siks. For this purpose, experimentally-deduced Sik target sequences were aligned and compared. Since only a few Sik targets are experimentally confirmed in *Drosophila*, we made use of proven targets from different organisms. This gave us a more generic, but a more conserved sequence of amino acids. SCARB1 and SREBP-1 were excluded from this analysis, since they do not contain any of the conserved residues. It is therefore possible that Siks have a second selection mechanism for phosphorylation. In particular, a 15 amino acid long region flanking the serine / threonine residue that is phosphorylated by Sik in different species was aligned to search for a target motif (**Figure 4A**). An amino acid sequence with 80% consensus was accepted as the target motif: [L/W]X[R/K]XX[S/T]*XXXL (* marks the serine / threonine residue phosphorylated by Sik) (**Figure 4B**). After determining the target motif, we scanned the *Drosophila melanogaster* proteome for proteins carrying this motif. In the whole fly proteome, this motif was identified in 307 annotated proteins, which include Notch (on serine 1066), Delta (on serine 740), and Fringe (on threonine 303), along with previously identified and confirmed targets (**Figure 4C, Supp. Table 3**).

**Figure 4.**
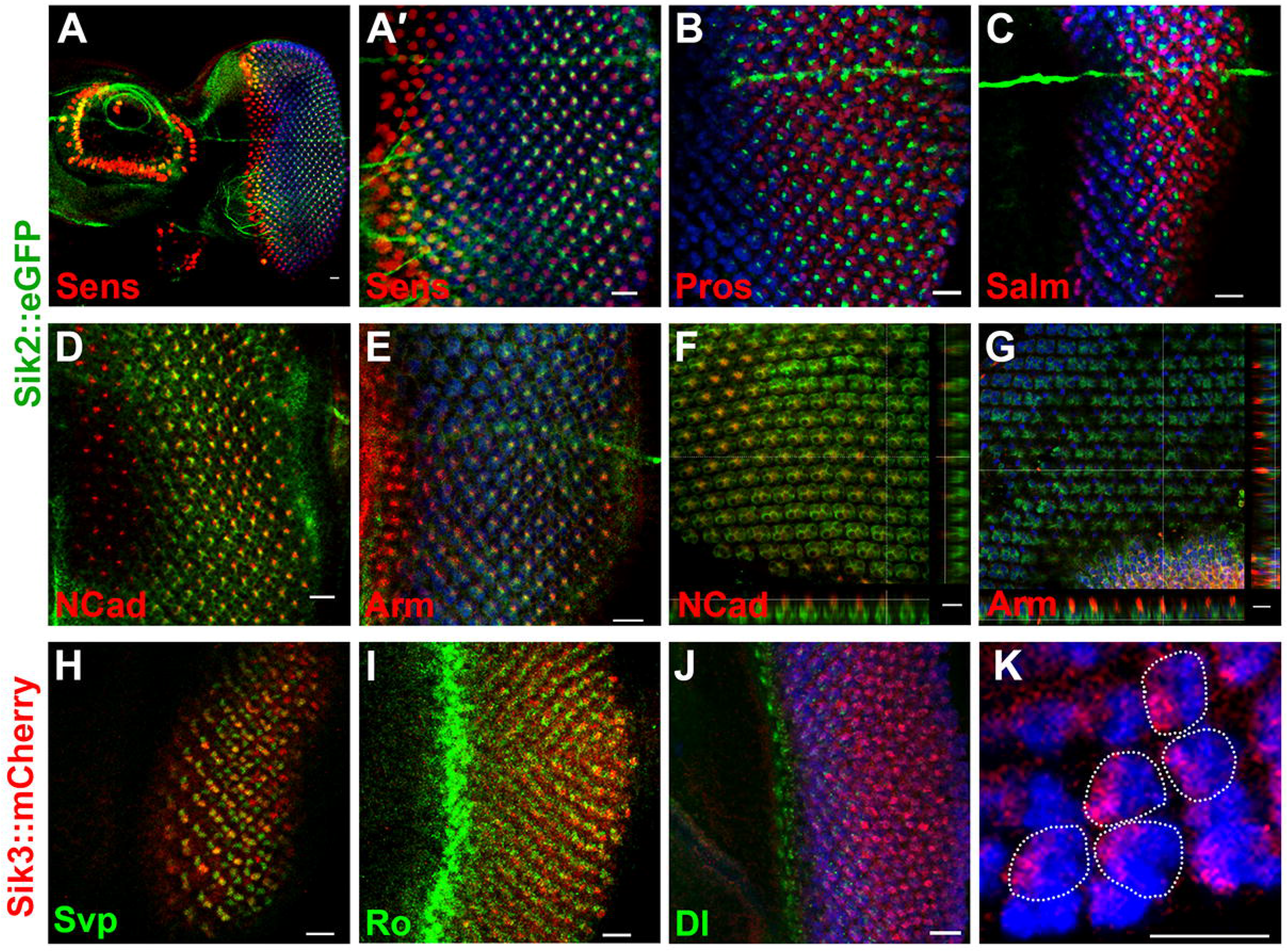
Salt inducible kinase target sequences. **A)** Protein sequences experimentally shown to be phosphorylated by Sik homologs were listed and aligned. 15 residues spanning the phosphorylated amino acid are shown in the figure. The *Homo sapiens* proteins start with letter “h”, *mus musculus* proteins start with letter “m”, *Drosophila melanogaster* proteins start with letter “d”. Sakamototide is a synthetic peptide based on CRTC2. The number of the residue to be phosphorylated by Sik is written after the underscore. “S” stands for serine, “T” stands for threonine. **B)** The motif was determined by 80% likelihood consensus sequence: [L/W]X[R/K]XX[S/T]*XXXL (* marks the phosphorylated serine / threonine residue). **C)** The motif was scanned against *Drosophila melanogaster* proteome. Notch, Delta and Serrate proteins comprise the consensus sequence to be phosphorylated by Sik. The serine (S) or threonine (T) residue to be phosphorylated is highlighted with turquoise and counted as 0. The positively charged amino acids (R, K) at −3 are colored red. Leucine (L) and Tryptophan (W) residues at −5 and +4 are colored green.

We reasoned that Siks might be modulating Notch pathway activity by protein phosphorylation. However, further experiments are necessary to verify the direct phosphorylation of these proteins by Sik. Unfortunately, we could not perform these experiments due to lack of suitable antibodies.

### Siks are Expressed in the Developing Retina

The observed phenotypes suggest a role for Siks in eye development and modulation of Notch pathway activity. This implies that Siks must be expressed in the developing retina. Thus, we set out to determine the localization of these proteins. Due to a high sequence conservation of Sik family members, most antibodies have the problem of cross-reactivity. To confirm their specificity, several commercially available antibodies (see Materials and Methods) were first tested in Western Blot (WB) analyses on crude fly extracts and immunofluorescence (IF) stainings on dissected larval and sectioned adult tissues. None of the commercially available α-SIK2 antibodies were able to specifically detect fly Sik2 in WB or IF (**data not shown**). Sik3 has two known isoforms. The long isoform has previously been shown to interact with the Salvador (Sav) protein (Wehr et al 2012). However, using the Abcam ab88495 α-human SIK3 antibody in WB, we observed predominantly the Sik3 short isoform. This antibody recognizes fly Sik3 at >75 kD (**Supp. Figure 2B**), slightly higher than the calculated size (77 kD), which was also confirmed by overexpression of Sik3 (**data not shown**). While we were able to show that *Drosophila* Sik3 is highly expressed in the developing larval nervous system in WB analyses, (**Supp. Figure 2B**), no specific staining for Sik3 could be obtained in IF analyses using the same antibody (**data not shown**).

The results we obtained using larval and adult tissues showed that none of the tested antibodies were specific enough to recognize and distinguish fly Sik2 or Sik3 in IF experiments. Thus, in order to observe endogenous protein localization, we generated BAC transgenic constructs, in which Sik2 and Sik3 were tagged with fluorescent proteins. BAC clones spanning the entire coding sequence, UTRs and 14 and 6 kb upstream regulatory region of the *Sik2* and *Sik3* genes respectively were used; green fluorescent protein (to Sik2) or mCherry (to Sik3) sequences were inserted C-terminally, prior to the stop codon using the P[acman] recombineering technique (Warming et al 2005) (**Supp. Figure 2A**) (see Materials and Methods).

Eye imaginal discs of BAC transgenic flies were co-stained for GFP or mCherry and for cell-type specific markers to identify the localization of Sik2 and Sik3 proteins, respectively. Senseless (Sens) was used as a marker for PR R8 (Frankfort et al 2001), Prospero (Pros) as a marker for R7 and cone cells (Cook et al 2003), Spalt-major (Salm) as a marker for R3/R4 cells for the initial few rows posterior to the MF and R7/R8 for the more posterior cells (Domingos et al 2004), Seven-up (Svp) was used as a marker for R3 and R4 (expressed stronger in R4 than in R3) and R1 and R6 (Fanto et al 1998), and Rough (ro) as a marker for PR R2/R5 (Frankfort et al 2001). Additionally, N-Cadherin (N-Cad) and Armadillo (Arm) were used to mark the Adherens Junctions (AJs). They are known to localize to the apical surface, between cone cells and PRs in pupal retinae (Izaddoost et al 2002) (Mirkovic & Mlodzik 2006).

In the late 3^rd^ instar larval eye discs, Sik2::eGFP signal was observed in the center of every ommatidium, starting 6-7 rows posterior to the MF (**Figure 5A-E**). The localization does not appear to be nuclear, thus no co-localization was observed with Sens (**Figure 5A, A′**), Pros (**Figure 5B**), or Salm (**Figure 5C**), all central photoreceptor markers that are localized to the nucleus (Frankfort et al 2001) (Cook et al 2003) (Domingos et al 2004). Sik2::eGFP was observed to be close to the N-Cad and Arm signals at the larval stage (**Figure 5D, E**), though the signals did not co-localize. At ~30% pupal development, the Sik2 signal was localized to PRs, in close proximity to Arm and N-Cad, but more basal and dispersed in the cytoplasm (**Figure 5F, G**).

**Figure 5.**
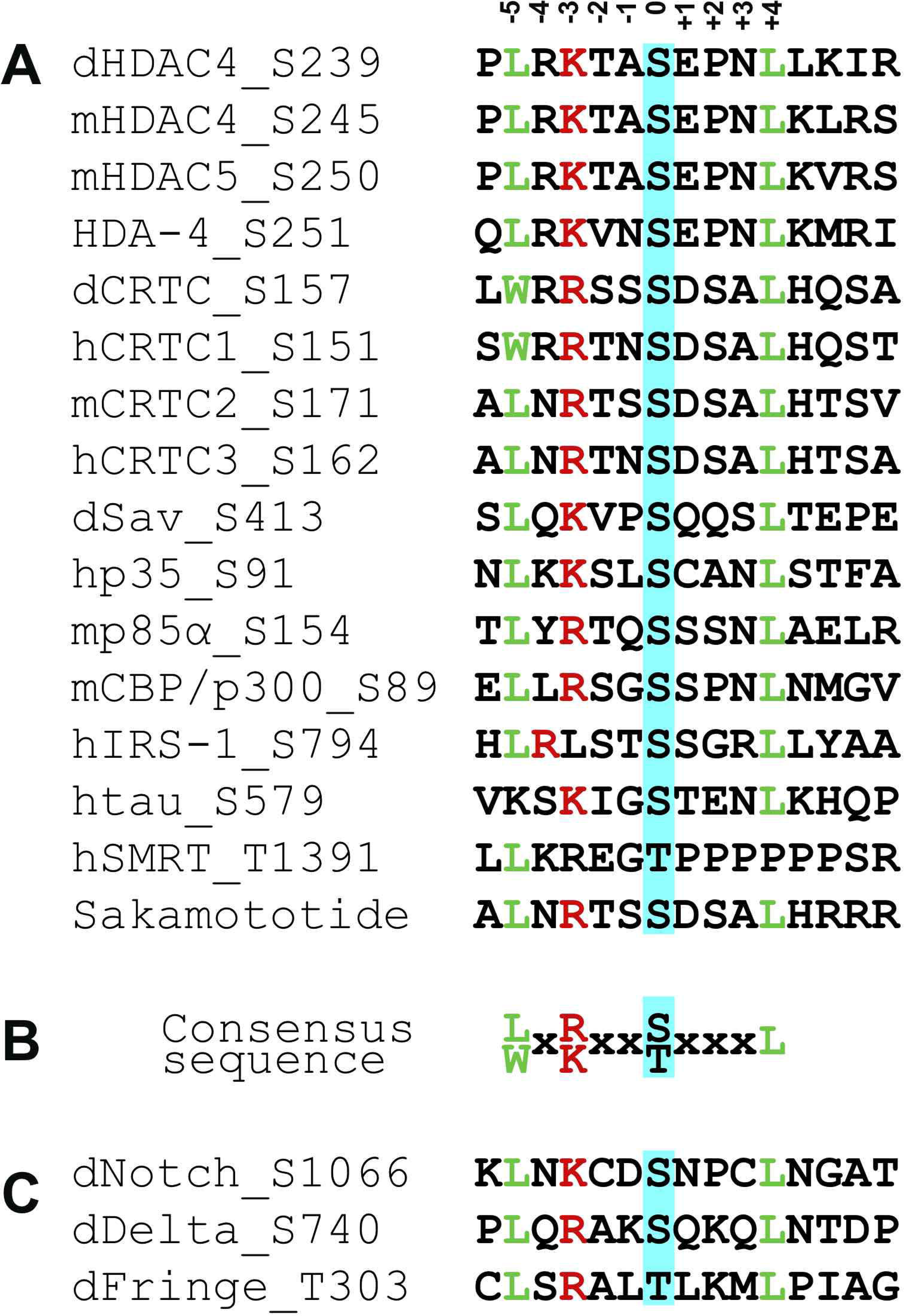
Expression profile of Sik2 and Sik3 in the developing retina. **A**, **A**′ and **J** are maximum projection. Rest of the acquisitions are single section, with ~0.5 μm thickness. **A, A**′**, B, C, D, E, H, I, J, K** are 3^rd^ instar larval eye disc. **F, G** are ~30% pupal eye disc. Wherever used, elav appears in blue, marking neural cells. **A-G**) Sik2::eGFP construct transgenics. Sik2 is shown in green **A**-**A**′) Senseless (Sens) shown in red, marking the photoreceptor R8. **A**′) Inset of A. **B**) Prospero (Pros) shown in red, marking R7 and cone cells. **C**) Spalt-major (Salm) shown in red, marking R3 and R4 in ommatidia immediately posterior to the morphogenetic furrow (MF) and R7 and R8 in more posterior ommatidia. **D**) N-Cadherin (N-Cad) shown in red, marking Adherens Junctions (AJs) at larval stage. **E**) Armadillo (Arm) shown in red, marking AJs at larval stage. **F**) N-Cad shown in red, marking AJs at pupal stage. **G**) Arm shown in red, marking AJs at pupal stage. **H-K**) Sik3::mCherry construct transgenics. Sik3 shown in red **H**) Seven-up (Svp) shown in green, marking R3/R4 in early ommatidia and R1/R6 in ommatidia more posterior. Sample is mid-3^rd^ instar larva. **I**) Rough (ro) shown in green, marking R2/R5 immediately posterior to the MF. **J**) Delta (Dl), the neurogenic factor shown in green. **K**) Inset of J. Dl signal is not represented here. The ommatidium borders are marked with dashed line. Posterior to the right and scale bar is 10 μm in all stainings.

Co-staining of Sik3::mCherry with Svp showed a co-localization with photoreceptors R3/R4 starting at row 3 (**Figure 5H**), and not with ro (**Figure 5I**). Delta (Dl) was expressed both in Sik3-positive and Sik3-negative cells in the initial few rows of ommatidia (**Figure 5J**); posterior to this, Sik3 was expressed in most PR cells (**Figure 5K**).

Taken together, Sik2 appears as a cytoplasmic, PR-specific protein, localizing close to cell junctions, and Sik3 as an R3/R4-specific protein initially and that is later expressed pan-neurally during larval PR development.

## Discussion

Precise regulation of signaling pathways and the interactions between the signaling proteins is required for proper tissue development. One of the fundamental aims of developmental biology is to identify the genetic interaction network that directs the development of a tissue. Although we are far from understanding the complete picture, dissecting the role of major interactors will get us closer to the answer. In this study, we focused on fly homologs Sik2 and Sik3, their role in development of the retina, and their involvement in tumor progression. *Sik* genes are members of the AMPK family that are conserved in animals and plants. They function as regulators of sodium sensing, cAMP signaling, metabolism, bone formation, immune cell maturation and neuroprotection (reviewed in (Wein et al 2018)). Although some Sik functions like gluconeogenesis and lipogenesis have been thoroughly studied, their involvement in neural development is quite limited.

Molecular analyses revealed that development and tumor progression actually share many aspects; there are fundamental molecular mechanisms common to both processes (Bossuyt et al 2009). *Drosophila* eyes have been used as model to investigate developmental processes for decades. Lately, they are also employed in tumorigenesis studies. The eye undergoing metamorphosis is a great tool to study both the genetic interactions that control *de novo* tissue generation, and the molecular mechanisms that control cell division to establish proper tissue size. With this motivation, we tested Sik behavior in an insect tumor model.

Although well-known for its developmental role, Notch pathway activity was shown to be disturbed in several cancer types (reviewed in (Nowell & Radtke 2017) (Aster et al 2017)). Our first experiment was a gain of function of the Notch receptor specifically in the eye, causing its slight overgrowth and sensitizing it for tumorigenesis. While the phenotype was less obvious in the “eyeful” background, in the “sensitized” background changes in Sik2 levels clearly induce tumor-like structures (**Figure 1**, **Supp. Table 1**). The misregulation of Sik2 and Sik3 in several cancer cases (Miranda et al 2016) (Amara et al 2017) at both genetic and epigenetic level (Fan et al 2019), and Sik3 being established as an ovary cancer marker (Charoenfuprasert et al 2011) supports this finding. In addition to the previous correlation studies, in this experiment we show that Sik misregulation can be a causative for tumor formation. Siks have been associated with certain cancer types, with both oncogenic and tumor suppressor roles, suggesting Sik involvement at a very fundamental level in cancer-related mechanisms. This dual role is also in parallel with Notch-mediated neoplasias where Notch acts either as oncogene or tumor suppressor gene depending on the cellular context (Bolos et al 2007).

Development is hand in hand with specification. The involvement of Siks in animal development has been shown in this study and in a few other reports. Thus, we suggest that Siks are at the intersection of these two pathways (Wehr et al 2012).

Among different experiments, although misregulation of Siks resulted in severe defects, we did not observe any phenotype with the simple knock-out of *Sik2* or *Sik3* (**data not shown, Supp. Figure 3**). Furthermore, the behavior of constitutively active Sik3 (*Sik3^S563A^*) resembled that of Sik2 (**Figure 1H**, **Figure 3E,L,O**, **Supp. Figure 4E**), suggesting a compensation mechanism between Sik paralogs, or with other members of the AMPK family. The overexpression of Sik proteins harboring point mutations that alter protein function are useful tools to escape a possible redundancy. Thus, we consider the results obtained by overexpression of constitutively active and kinase-dead versions of Siks to be more valid, and we actually observed stronger phenotypes with these transgenes than with simple knock-down experiments.

As demonstrated in **Figure 1**, Siks contribute to tumorigenesis and metastasis. It is known that reduction in E-Cadherin and loosening AJs is key for the epithelial-mesenchymal transition and consequently metastasis (Goodwin et al 2014) (Yao et al 2016), and a compromise in AJ stability is correlated with the invasive phenotype. Sik2 observed at the cell junctions (**Figure 5D-G**) is in line with this finding. We have previously shown that Sik alters invasiveness of the tumor cells (Zohrap et al 2018). Other reports propose Siks as anti-metastatic proteins (Bailey et al 2009) (Xia et al 2018) and show that loss of Sik function disables anoikis (Cheng et al 2009). Combining the invasiveness phenotype with the effect of Sik on survival, Siks are strong candidates for promoting cancer and anoikis.

The ultrastructural analysis on the gain of function and loss of function alleles of Sik2 and Sik3 revealed that, correct levels of Sik proteins are required for proper development of the eye. Any misregulation of Sik expression results in disruption of the adult lens and retinal bristle structure, independent of increase or decrease of Sik function (**Figure 2A-B′′, E-F′′**). Interestingly, misexpression of transgenic constructs carrying Sik point mutations not only disturbed the cellular organization, but also the eye morphology. The constitutively active and kinase-dead mutants lead to a “ventral notch” in the eye (**Figure 2G-H′′**). Comparison with the control eye, it appears more like a growth defect specific to ventral half of the eye, rather than an extension in dorsoventral axis. This indicates a D-V polarization problem that highly resembles Notch pathway phenotypes (Chern & Choi 2002) (reviewed in (Singh et al 2012)). Presence of flies missing the entire ventral eye due to strong knock-down of Sik2 (**Figure 2D**) further increases the possibility that Siks function at the L2 stage in dorsoventral polarization of the developing eye tissue, in a Notch-dependent manner and that particularly Sik2 interacts with the Notch pathway or its effectors. The observation of eye-to-antenna homeotic transformations (**Figure 1E**, **Figure 2C**′′) is yet another support to this hypothesis, relating to the fact that eye and antenna are specified from a single continuous epithelium via Notch and EGFR expression during early L2 stage (**Supp. Figure 1A**) (Kumar & Moses 2001). The forked bristle phenotype observed in Sik2 overexpression and Sik point mutation backgrounds resembles known Notch phenotypes in the mechanosensory system (**Supp. Figure 4**). As Notch is employed several times during neural development, we suggest that Siks might be affecting the activity of the Notch pathway and are involved at different steps of eye development. However, the exact contribution of Siks in this process needs still to be established.

To test if Siks interfere with the Notch pathway, we analyzed their genetic interaction. We found that manipulation of Sik2 levels exaggerate the small eye phenotype of Serrate knock-down (**Figure 3G-J**) and of Fringe overexpression (**Figure 3M,N**), although they do not affect eye size on their own (**Figure 2B-B′′, G-H′′**). Unlike Fringe and Serrate, Delta overexpression increases eye size, since it is not only required for establishment of D-V polarity at the L2 stage, but also for neurogenesis behind the MF in the L3 stage (Baonza & Freeman 2001). Similarly, changes in Sik2 levels exaggerate or modify this phenotype (**Figure 3A-C, F**). Although we have not verified a direct interaction between Siks and Notch pathway members, they are epistatic to each other and our data clearly show that Sik2 can modify the Notch pathway output. Particularly, while Sik2 knock-down strongly suppresses eye growth (**Figure 2C**), Delta overexpression can rescue this phenotype (**Figure 3F**). The ability of Delta to suppress the Sik2 phenotype suggests that Delta acts downstream of Sik2. On the other hand, Sik3 overexpression or knock-down did not modify the original phenotype (**Figure 3 D, K**, **data not shown**), suggesting that the genetic interaction is mostly through the Sik2 protein. As in the first experiment (**Figure 1**), the constitutively active allele of Sik3 rather behaves like Sik2 (**Figure 3E,L,O**) and modifies the Notch pathway member phenotypes.

After confirming a genetic interaction between Sik and Notch, we asked if Notch pathway members can be direct targets of Siks, since they are functional kinase enzymes and phosphorylation is a key regulation mechanism in signal transduction. By comparing the sequences of experimentally proven Sik targets, we identified a consensus phosphorylation motif (**Figure 4A-B**). Analysis of the fly proteome showed that >300 reviewed protein entries contain this deduced motif; the list includes Notch, Delta, Fringe and UVRAG from the Notch pathway (Lee et al 2011). We suggest that Siks can target one or more Notch pathway members and through phosphorylation alter the signaling outcome. The complex phenotype we observed may even be due to several related targets being phosphorylated by Siks. These putative phosphorylation sites need of course to be confirmed biochemically, once specific antibodies become available.

Scanning the fly proteome revealed experimentally-confirmed Sik targets like Salvador, increasing the validity of our screening. Further hits contain several DNA and RNA binding proteins, transcription coactivators, proteins important for the RNAi machinery, epigenetic regulators, some proteins important in cell cycle and cancer, proteins related to cytoskeletal organization, proteins functioning in signal transduction, some proteins functioning in phototransduction, gustatory and odorant receptors, several ubiquitin-protein ligases, some proteins important in lipid synthesis, many mitochondrial proteins, and few proteins with metallopeptidase activity, important for migration. Our data hint to several proteins related to animal development and retina morphogenesis that could be Sik targets, including Pannier, Slingshot, tiptop, hedgehog, hopscotch, Calpain-D, Neurobeachin, fork head, prospero, FGFR2, flamingo, elav, Dscam2, Patj, scribble, arrowhead and domeless (**Supp. Table 3**). Pannier is particularly interesting, since it is responsible for establishing the dorsal eye fate and we show that Sik disruption leads to a ventral eye defect.

Recent findings point to a potential role of these kinases in animal development, mostly at the skeletal tissue. We aimed to investigate the role of salt inducible kinases in development, with the caveat of lack of specific fly antibodies. With the only working α-SIK3 antibody, our WB analyses showed that Sik3 was highly expressed in the larval nervous system (brain and imaginal discs) (**Supp. Figure 2B**), unlike the ubiquitous expression shown previously (Katoh et al 2004). In contrast to a previous report (Wehr et al 2012), we mainly detected the short alternative transcript of Sik3 (**Supp. Figure 2B**) and thus focused on the short Sik3 transcript to generate an overexpression construct (**Supp. Figure 2C**) and for genomic tagging with a fluorescent protein.

To overcome the antibody problem, we generated novel useful tools. Using the BAC recombineering technique, we created translational fusion proteins of Siks with fluorescent protein tags (**Supp. Figure 2A**). IF analyses of retinae of the Sik fusion protein-expressing animals showed that they are expressed in a specific pattern, for a limited period in development. In our analyses, Sik3 expression was detected only in late L3 stage, in R3/R4 for the first few rows after the MF, and many other neural cells in the more posterior compartment. The stepwise expression of Sik3 in PRs suggests a potential role of Sik3 in PR specification. We detected Sik2 expression adjacent to the AJs in late L3 stage, and in the cytoplasm of PRs, beneath the AJs at the pupal stage. This localization can be particularly meaningful. A previous report showed that AJs are regulated partly by Sik3 and Lkb-1, the major regulator of Siks. In the mutants of Lkb1 and Sik3, AJs are defective and Armadillo is mislocalized (Amin et al 2009). These clues suggest that, in addition to cell cycle control and cell death, Sik2 might function in development by regulating cell-to-cell interactions, cell morphology and polarity, similar to Sik3. Cytoplasmic localization of Sik2 also suggests that its nuclear targets such as HDACs (Wang et al 2011) and CRTCs (Choi et al 2011) can be excluded in the pupal stage.

Taken together, this study shows that fly Sik2 and Sik3 are required for proper retinal development, and they function through the Notch pathway. Further studies will uncover if their interaction is direct, and at exactly which stage it takes place.

## Materials and Methods

### Fly genetics and Husbandry

Flies were raised at 25 °C, on regular fly food.

*Sik2*^*Δ41*^, *UAS-Sik2-myc* and *UAS-Sik2-K170M* lines were kindly provided by Jongkyeong Chung (Korea) (Choi et al 2011). *Sik3*^*Δ109*^, *neoFRT42D, Sik3*^*Δ109*^, *UAS-Sik2-S1032A* and *UAS-Sik3-S563A* were a kind gift from Nic Tapon (UK). Sensitized and eyeful lines were kindly provided by Bassem Hassan (Paris) (Bossuyt et al 2009). *UAS-Dicer; ey-GAL4, lGMR-Gal4 / CyO* were a kind gift from Claude Desplan (New York).

The following fly lines were obtained from the Bloomington Drosophila Stock Center: *w^1118^* (3605), *UAS-Notch* (26820), *UAS-Delta* (5612), *UAS-Ser* (5815), *UAS-Fringe* (5831), *UAS-Delta.DN* (26697), *lGMR-GAL4* (8121), *ey-GAL4* (5535), *elav-GAL4* (458), *mirr-GAL4* (29650), *sca-GAL4* (6479), *repo-GAL4* (7415), *neoFRT42D, GMR-myr-GFP* (7110), *neoFRT42D, GMR-hid / CyO; ey-GAL4, UAS-FLP* (5251). *actin-GAL4* (25374). Following RNAi lines were supplied by the VDRC Stock Center: *Sik3-RNAi* (107458), *Sik2-RNAi* (103739), *Ser-RNAi* (108348), *Delta-RNAi* (3720), *Fringe-RNAi* (51977).

### Generation of transgenic fly lines

The *UAS-Sik3::T2A::mCherry* construct was generated with the full-length cDNA of *Sik3* isoform A. The pUAST-attB vector was modified to include the T2A::mCherry fusion protein. The T2A::mCherry plasmid was a kind gift from Dr. Stefan Fuss (İstanbul). It was cloned into the pUAST-attB cloning site using the restriction enzymes *Xho*I and *Kpn*I. The cDNA template FBcl0162999, corresponding to *Sik3*, was purchased from Drosophila Genomics Resource Center (DGRC). *Sik3* cDNA was amplified by high fidelity DNA polymerase Advantage 2 (Clontech) and cloned into pUAST-attB-T2A::mCherry using the *Not*I and *Xho*I restriction sites, T2A and mCherry being N-terminal. Transgenic fly lines were generated at Genetivision, Inc.

### Recombinase mediated recombineering

Clones 149A06 for *Sik2* and 12M10 for *Sik3* genes were ordered from Berkeley Drosophila Genome Project (BDGP) in attB-P[acman]-Cm^R^-BW BAC vectors. The clones include the entire coding sequence, untranslated regions, 14 kb and 6 kb upstream regulatory sequence for *Sik2* and *Sik3* genes respectively. Isoform A (CG42856-RA / NM_137515) was targeted for the *Sik3* gene; *Sik2* has no alternative transcripts. The clones were transformed into modified SW102 strain of *E. coli*, which is missing the *galactokinase* gene (*galK*) from the galactose operon (Warming et al 2005). PCR primers were designed with large homology arms to target immediately before the stop codon of the *Sik* genes. *galK* gene was amplified by PCR using these primers. The PCR products were transformed into the cells and the fragments were introduced to *Sik* genes by homologous recombination. *Recombinase* was induced by heat shock at 42°C. The bacteria were incubated on minimal media and on MacConkey indicator plates to select for *galK*. Next, the targeted fluorescent genes were amplified by PCR using primers with homology arms. The *galK* gene was replaced with the fluorescent gene by recombineering and the colonies were counter-selected on minimal media including D-galactose and DOG indicator. The clones were verified by restriction digestion and sequencing. The translational fusion of Sik and the fluorescent tags were generated. During the PCR primer design, a 3 amino acid linker (Gly-Ser-Gly) was inserted between Sik and the fluorescent genes to allow independent folding of the proteins (Warming et al 2005). Transgenic flies were generated by GenetiVision, Inc.

### Generation of tissue specific clones

Entire eye clones were obtained using the *GMR-hid* construct, which induces apoptosis in the tissue even when heterozygous. A copy of *neoFRT42D, GMR-hid* was replaced with *neoFRT42D, Sik3*^*Δ109*^ in mitosis dependent recombination, by Flippase expressed by *ey-GAL4* driver.

Regional, mitosis dependent clones were obtained using the MARCM technique. A copy of *neoFRT42D, GMR-myr-GFP* was replaced with *neoFRT42D, Sik3*^*Δ109*^, by Flippase expressed from the eyeless promoter. *Sik3* null mutant cells were recognized by loss of red eye pigmentation.

### Immunohistochemistry

Fly tissues were dissected in cold PBS and fixed in 4% formaldehyde, at room temperature for 20 minutes. After 3 washes in PBX (0.3% TritonX-100 in 1X PBS) the tissues were blocked for 1 hour at room temperature in serum added BNT (10% donkey serum, 0.01% BSA, 250 mM NaCl, 0.1% Tween-20, PBS 1X). Tissues were incubated with primary antibodies overnight at 4°C. After 3 washes in PBX, the tissues were incubated with secondary antibodies for 2 hours at room temperature. After final washing steps, tissues were mounted in Vectashield^©^ mounting medium.

Antibodies were used at following concentrations: rabbit α-SIK3 (Abcam ab88495, 1:50 for IHC, 1:500 for WB), goat α-SIK2 (Santa Cruz sc-33074, 1:200 for WB), mouse α-actin (Cell Signaling 8H10D10, 1:1000 for WB), mouse α-myc (Cell Signaling 9B11, 1:1000 for WB), rabbit α-DsRed (Clontech 632496, 1:500 for WB, 1:100 for IHC), chicken α-GFP (Abcam ab13970, 1:1500 for IHC), Guinea pig α-Senseless (kindly provided by H. Bellen, 1:500 for IHC), rabbit α-Spalt (Salm) (1:150 for IHC, kindly provided by F. Schnorrer), rat α-DNCad (DSHB DN-Ex#8, 1:20 for IHC), mouse α-Armadillo (DSHB N2-7A1, 1:50 for IHC), mouse α-Seven-up (kindly provided by Y. Hiromi, Japan, 1:500 for IHC), mouse α-rough (DSHB 62C2A8, 1:50 for IHC), mouse α-Prospero (DSHB MR1A, 1:5 for IHC), mouse α-Delta (DSHB C594.9B, 1:50 for IHC), rat α-elav (DSHB 7E8A10, 1:20 for IHC), mouse α-repo (DSHB 8D12, 1:20 for IHC).

### Imaging / Electron Microscopy

Immunohistochemistry (IHC) sample images were acquired using a Confocal TCS SP5 system. The whole mount fly images (white light and fluorescent) were acquired with a Leica DFC310 FX system, after samples were mounted in pure ethanol.

Whole mount detailed fly images were acquired with SEM (Scanning Electron Microscopy), by Boğaziçi University Advanced Technologies Research and Development Center, Electron Microscopy Facility. Gold coating was not applied.

### Protein analyses

Crude protein was extracted from dissected fly tissues in following buffer: 50 mM NaCl, 50 mM Tris-Cl pH 7.5, 10% Glycerol, 320 mM Sucrose, 1% Triton-X100, Complete Protease Inhibitor Cocktail^©^ Inhibitor (Roche). The tissues were homogenized and centrifuged for 10 minutes at 13,200 rpm, 4°C. The crude protein was loaded on an 8% SDS-PAGE, after denaturation by DTT. Separated proteins were transferred to a PVDF membrane with a pore size of 0.45 μm. The membrane was blocked in 5% non-fat milk in TBS-T (20 mM Tris-Cl pH 7.6, 150 mM NaCl, 0.1% Tween-20) and blotted with primary antibody in 5% milk solution, overnight at 4°C. Signals were detected by LumiGLO^®^ Reagent (Cell Signaling).

### Bioinformatic analyses

The protein sequences were downloaded from UniProtKB (http://www.uniprot.org/) and Protein similarity was calculated by EBI global protein alignment tool, Needle (EMBOSS) (http://www.ebi.ac.uk/Tools/psa/emboss_needle/), by pairwise comparison of the protein sequences of interest.

To establish the recognition pattern for salt inducible kinases, known Sik target proteins were chosen manually from publications. Only the experimentally proven target sequences were selected. Targets of different Sik homologs were pooled together (UniprotKB numbers are given in parentheses): *Drosophila* HDAC4 (Q9VYF3) (Wang et al 2011), mouse HDAC4 (Q6NZM9) (Wang et al 2000), mouse HDAC5, (Q9Z2V6) (Takemori et al 2009), *C. elegans* HDA-4 (O17323) (van der Linden et al 2007), *Drosophila* CRTC (M9FPU2) (Choi et al 2011), human CRTC1 (Q6UUV9) (Katoh et al 2006), mouse CRTC2 (Q3U182) (Screaton et al 2004), human CRTC3 (Q6UUV7) (MacKenzie et al 2013), *Drosophila* Sav (Q9VCR6) (Wehr et al 2012), human p35 (Q15078) (Sakamaki et al 2014), human p85α (P27986) (Miranda et al 2016), mouse CBP/p300 (Q09472) (Bricambert et al 2010), human IRS-1 (P35568) (Horike et al 2003), human tau (P10636) (Yoshida & Goedert 2012), human SMRT (Q9Y618) (Qu et al 2016) and Sakamototide^®^ (synthetic peptide) (Berggreen et al 2012). Since *C. elegans* and *D. melanogaster* protein targets are missing, S/T at the −2 of the phosphorylation motif was ignored. The target sequences were aligned via EBI MView multiple alignment viewer using the default parameters (http://www.ebi.ac.uk/Tools/msa/mview/) (Brown et al 1998). Consensus sequence of 80% was accepted as the motif. The common pattern was searched against *Drosophila melanogaster* proteome, by ScanProsite tool of ExPaSy (http://prosite.expasy.org/scanprosite/), using default parameters. It returned 507 hits; 307 were reviewed UniProtKB entries. Known targets and novel candidates were analyzed in the results. Notch protein (P07207) was targeted on serine 1066, Delta protein (P10041) on serine 740, and Fringe protein (Q24342) on threonine 303.

## Acknowledgements

The authors declare no conflict of interest. This project was funded by EMBO installation grant 1656, BAP 10B01P12, and BAP5682. HBŞ was supported by a TÜBİTAK BİDEB 2218 postdoctoral fellowship. We thank Tiffany Cook for comments on the manuscript and Hugo Bellen, Sekyu Choi, Claude Desplan, Stefan Fuss, Bassem Hassan, Yash Hiromi, Frank Sprenger and Nic Tapon for the generous gifts of materials. We thank the Bloomington Drosophila Stock Center at the University of Indiana, Vienna Drosophila RNAi Center and Developmental Studies Hybridoma Bank at the University of Iowa for kindly providing the *Drosophila* stocks and the antibodies used in this study.

Data available on request from the authors.

## Author Contributions

Conceived and designed the experiments: HBŞ, SS, KB, AÇ. Performed the experiments: HBŞ, SS. Analyzed the data: HBŞ, SS. Contributed reagents/materials/analysis tools: KB. Wrote the paper: HBŞ, SS, KB, AÇ.

---

**Supplementary Figure 1. Larval development of *Drosophila* retina. A-C)** Schematic drawing of eye-antennal disc development. **A)** During the early second instar larval stage, posterior part of the epithelium is specified as eye precursor, via Notch receptor (shown in orange); anterior part is specified as antennae via EGFR. **B)** During late second instar stage, dorsal-ventral polarity is established. Notch receptor is expressed in the midline cells (orange). Notch is activated by Delta ligand on the dorsal half and by Serrate ligand on the ventral half. Fringe is expressed on the ventral half and suppresses the Notch activation by Serrate anywhere else than the dorsal-ventral border. **C)** During the late third instar stage, morphogenetic furrow (MF) sweeps the eye disc posterior to anterior and initiate retinal cells specification. Delta ligand is expressed behind the MF to induce neurogenesis. Anterior is to the left, ventral is down in all drawings. The drawings are not in-scale.

**Supplementary Figure 2. The Sik transgenic lines and mutant generated in this study. A**) The protein structure and genetic constructs of *Drosophila* salt inducible kinases. Sik2 and Sik3 protein kinase domains are shown in red, the UBA domains in green, and the key residues to suppression of Siks by PKA (S1032 and S563, respectively) are shown in blue. *Sik2::eGFP* and *Sik3::mCherry* are translational fusions, generated on BAC clones, where fluorescent proteins are added prior to the stop codon, after a flexible linker. Both clones comprise all the introns, exons, UTRs and sufficient regulatory region (14 kb and 6 kb for *Sik2* and *Sik3* respectively) to reflect the endogenous expression. UAS-Sik3::T2A::mCherry was generated using the EST clone encoding Sik3 CDS isoform A. mCherry was attached after a self-cleaving T2A linker. The drawings are in scale except for the regulatory region. **B**) Western blot on the wild type (*W^1118^*) strain crude protein extract from adult head, adult body, larval CNS (brain and eye-antennal imaginal discs) and larval body. >75 kD band was revealed with the α-human SIK3 antibody (estimated Sik3 size ~77 kD). Actin was used as loading control. **C**) Western blot of fly head extract, from control (*W^1118^*) and ectopic expression of Sik3 by eye-specific drivers *(ey-GAL4, lGMR-GAL4 / +; UAS-Sik3::T2A::mCherry / +*), revealed with α-DsRed antibody. Majority of the mCherry was already cleaved from the construct, as seen at >25 kD, and a small portion was still attached to Sik3, as seen at >100 kD (estimated size ~27 kD and 115 kD respectively).

**Supplementary Figure 3. Eye clones of Sik3 mutant.** Since *Sik3* null mutants are early stage lethal, eye-specific null mutant clones were generated. **A-A**′′) Full eye mutant for *Sik3* obtained by mitosis-dependent flippase, selected against *GMR-hid*. **A)** Control of the eye specific pro-apoptotic element *GMR-hid* allele (Control-hid). **A**′) Control of the *flippase* allele (Control-Flp). **A**′′) Full eye clone of Sik3 null mutant (*Sik3*^Δ*109*^). **B-B**′) MARCM for *Sik3* null mutant. **B)** Control of MARCM with *GFP* allele (MARCM Control). **B**′) MARCM clones (*Sik3*^Δ*109*^ MARCM). Red region is heterozygous with one copy of GFP and one copy of Sik3 null mutant (*Sik3*^Δ*109*^ / *GMR-myrGFP*). White region is the clone, homozygous for Sik3 null mutation (*Sik3*^Δ*109*^). The full genotypes are listed in **Supp. Table 2**. In all pictures, anterior is to the left, ventral is down.

**Supplementary Figure 4: Siks are important for bristle development. A**) Control of ubiquitous driver (*actin-GAL4 / CyO*) **B-E**) Sik modulation by different drivers causes reduction in size and defect in the shape of the macrochaetae. **B**) Ubiquitous ectopic expression of kinase-dead Sik2 (>*Sik2^K170M^*). **C**) Ectopic expression of constitutively-active Sik2 (>*Sik2^S1032A^*). **D**) Sik2 overexpression (>*Sik2* OE) in the bristles; leads to defect in microchaetae as well. **E**) Constitutively-active Sik3 (>*Sik3^S563A^*) expression in the macrochaetae. **F**) Delta dominant-negative (>*Delta^DN^*), causing supernumerary bristles. The full genotypes are listed in **Supp. Table 2**. Dorsal view in all pictures, posterior is down.

**Supplementary Figure 5. Siks are conserved in evolution.** Pairwise comparison of *Drosophila melanogaster* and *Homo sapiens* SIK2 and SIK3 proteins by global alignment. Kinase domains are highlighted with pink, the critical lysine residues in kinase domain (SIK2^K170^, SIK3^K70^) are highlighted in red, the Lkb-1 target in T-loop (SIK2^T296^, SIK3^T196^) are highlighted with yellow, the ubiquitin associated domains (UBA) were highlighted in green, the PKA target serine (SIK2^S1032A^, SIK3^S563A^) are highlighted in blue. Siks, especially the kinase domains are highly conserved in evolution. Human SIK2 and fly Sik2 kinase domains are 88.9% similar; human SIK3 and fly Sik3 domains are 85.3% similar; fly Sik2 and fly Sik3 domains are 82.5% similar.

**Supplementary Table 1. Eye phenotypes in sensitized and eyeful background.** The eye phenotype quantification in the sensitized (*Delta* overexpression) and eyeful backgrounds (*Delta* overexpression in combination with *GS88A8* epigenetic regulator mutation). Ratios for different backgrounds are listed and shown in the histogram. The not affected / normal eyes are similar to the parents’, which is slightly bigger than the wild type flies. The affected eyes are subclassified as folding / overgrowth (bigger eyes with at least one folding, or overgrowth of the eye), multiple eyes (ectopic eye tissue on the head surface or duplicate eyes on one side of the head), eye loss (total loss of the eye tissue, or eye-to-antenna differentiation), and metastasis (ectopic eye tissue in the body or in the head, which is not exposed on the surface). The incidences of normal eyes and affected eyes were quantified, and the ratio over the total number of eyes is presented in the table. The total ratio of affected eyes are shown in the last column for each background, together with the standard error of the mean obtained from 3 independent trials. The genotypes are as in **Figure 1**. The full genotypes are listed in **Supp. Table 2**.

**Supplementary Table 2.** The full genotypes used in this study.

**Supplementary Table 3. Potential Sik targets in***Drosophila* **proteome.** This data is supplied in the Excel table. Notch pathway members are marked with pink. Other interesting targets are marked with orange.

