## Supplementary figures and images for "Salt Inducible Kinases as Novel Notch Interactors in the Developing *Drosophila* Retina"

### Supplementary Figure 1

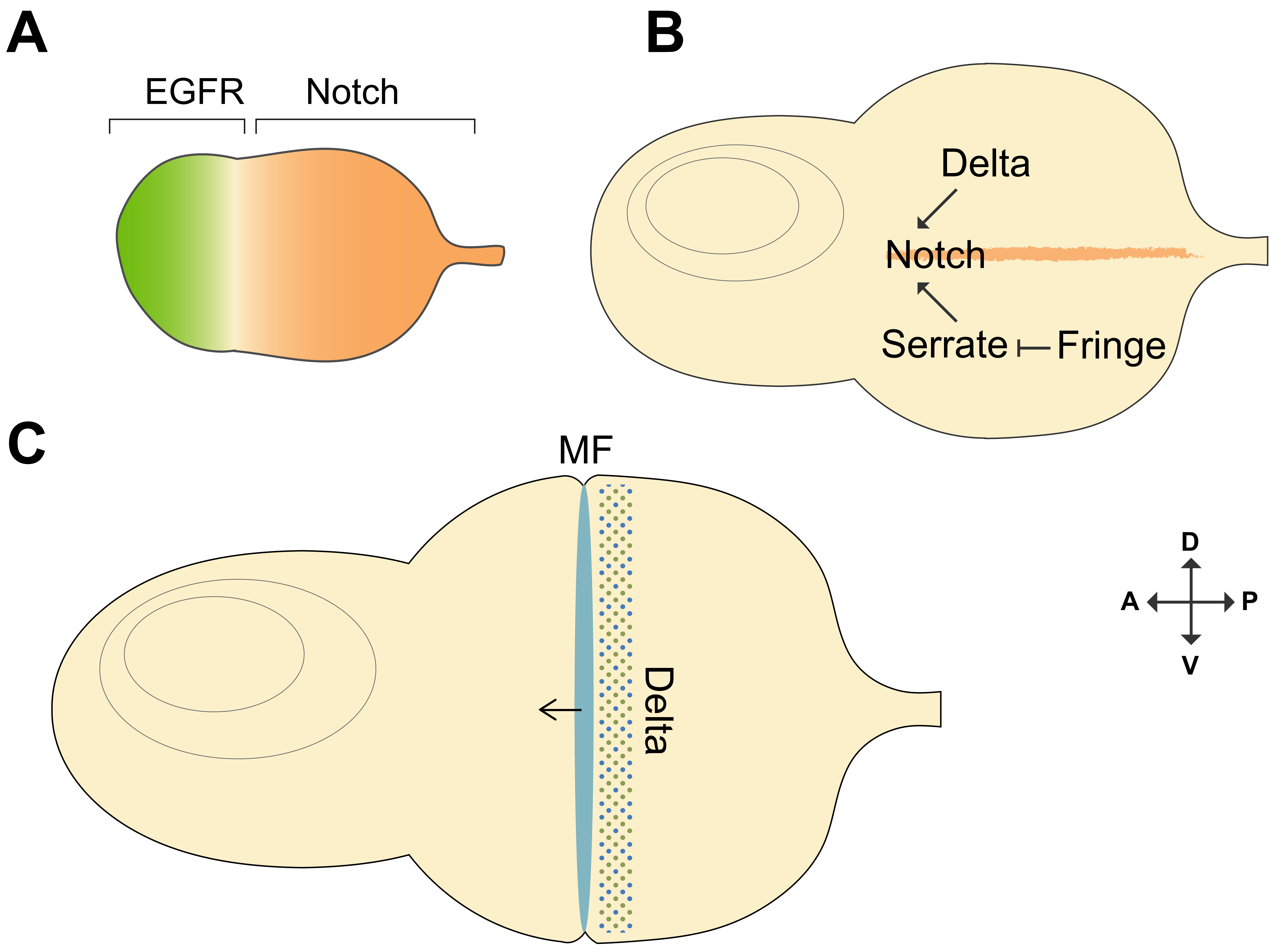

### Supplementary Figure 2

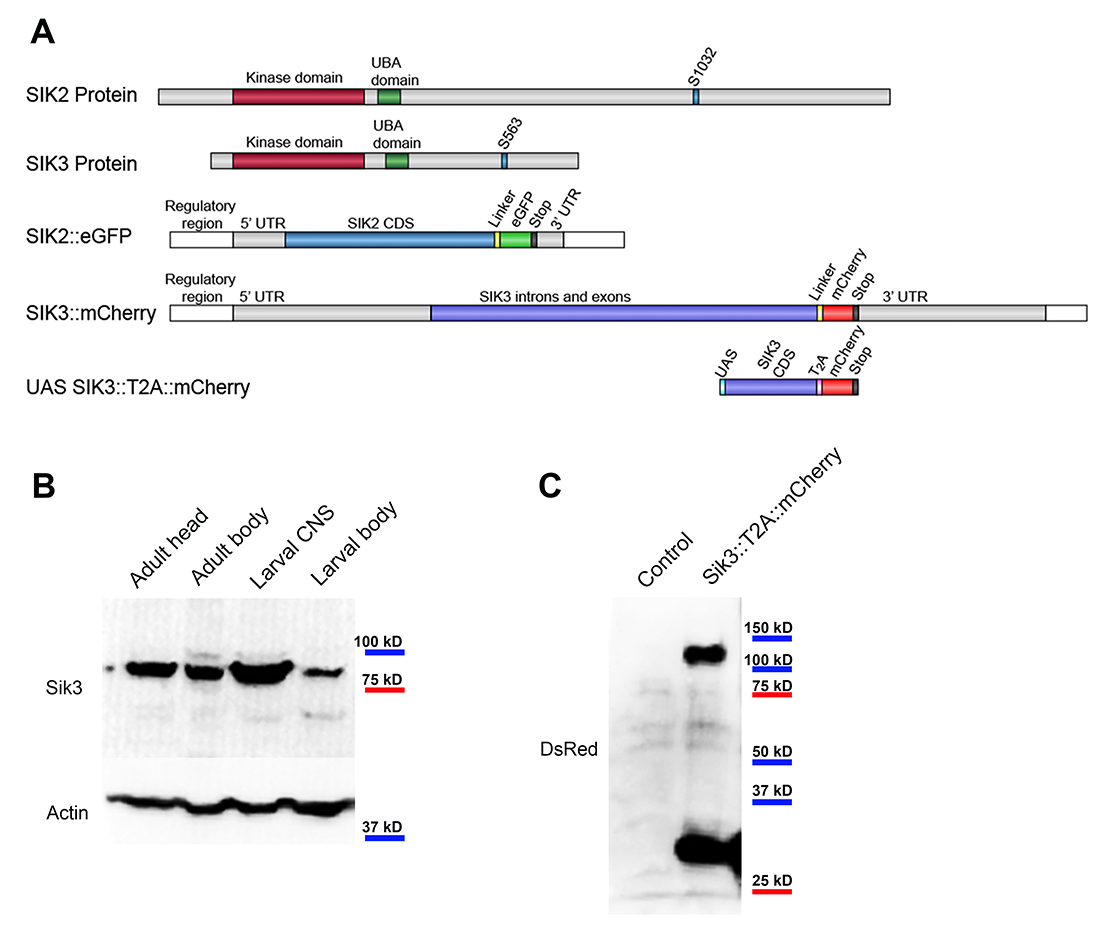

### Supplementary Figure 3

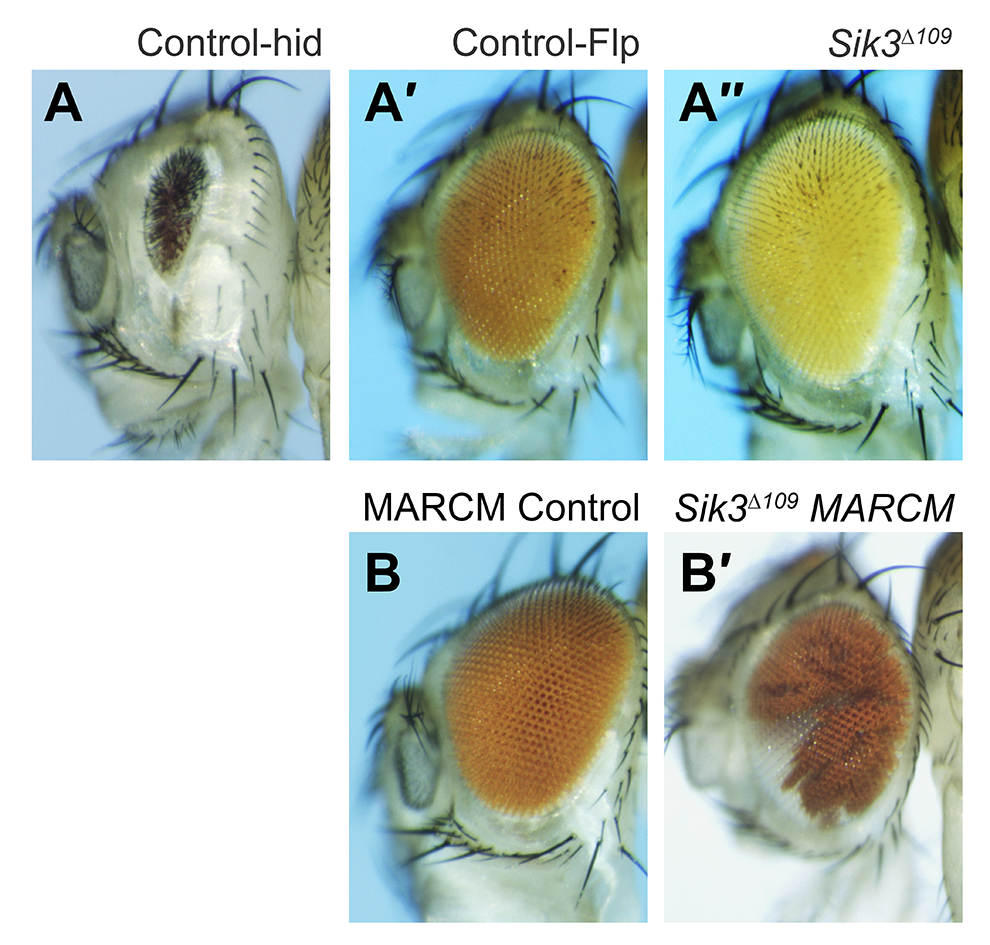

### Supplementary Figure 4

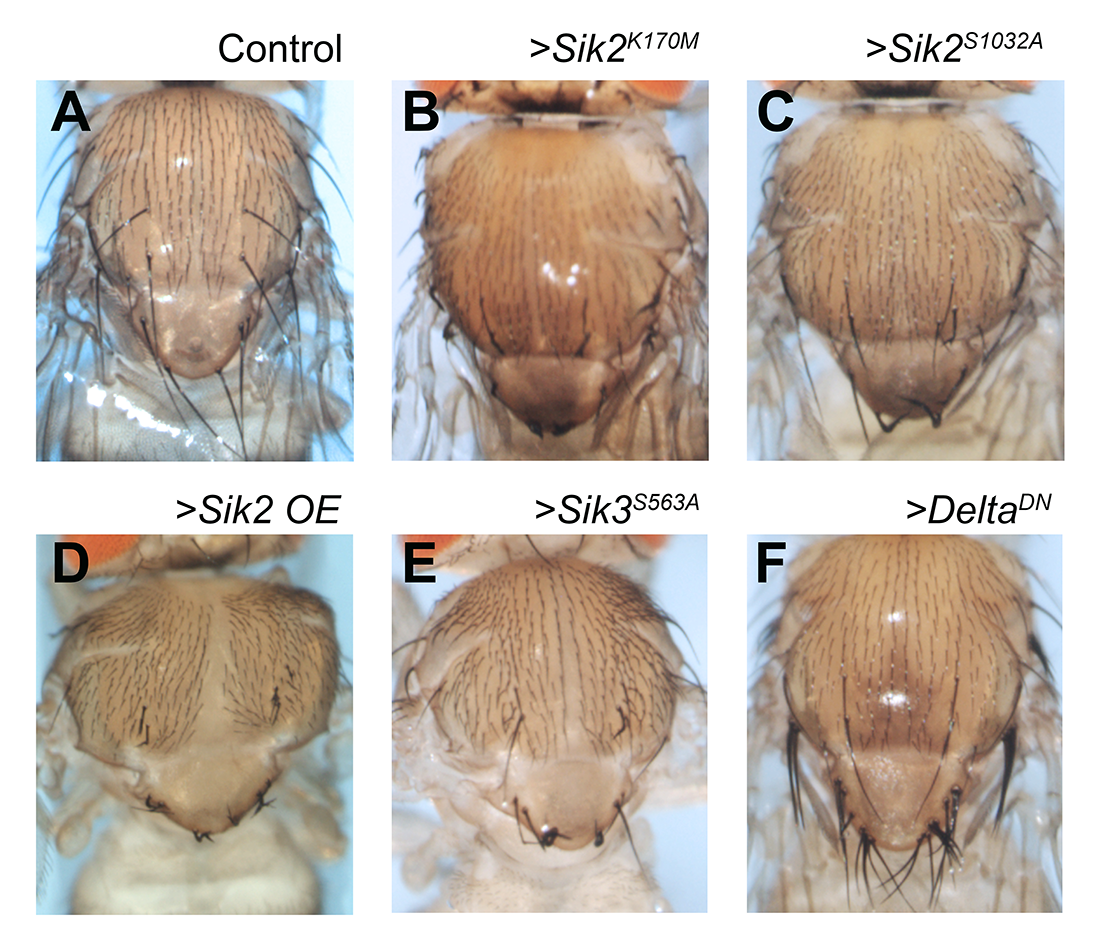
