## Supplementary Figure 5 for "Salt Inducible Kinases as Novel Notch Interactors in the Developing *Drosophila* Retina"

```
Fly_SIK2       1 MSTCEAAAAGENGSQAKSEESQPPEDQKKKQPQHERNEKQLDKPPENLPQNGKTEAKGAEGACSHPPLDALRSSVLLDAGAASPSIDAIVACKDALLAQK  100

Human_SIK2     1 ----------------------------------------------------------------------------------------------------  0

Fly_SIK2     101 LFASGGGSTPGPSPTSSAVGAGGISGKDLLKLKEPMRVGFYDIERTIGKGNFAVVKLARHRITKNEVAIKIIDKSQLDQTNLQKVYREVEIMKRLKHPHI  200
                                  .|.|.|.....    :.|:||||||.|:|||||||||||.||||||.|||||||||||||..||:|:||||:|||.|.||||
Human_SIK2     1 -----------------MVMADGPRHLQ----RGPVRVGFYDIEGTLGKGNFAVVKLGRHRITKTEVAIKIIDKSQLDAVNLEKIYREVQIMKMLDHPHI  79

Fly_SIK2     201 IKLYQVMETKNMIYIVSEYASQGEIFDYIAKYGRMSESAARFKFWQIISAVEYCHKKGIVHRDLKAENLLLDLNMNIKIADFGFSNHFKPGELLATWCGS  300
                 |||||||||:|:|:|:|||..||||||:|.:||::||.||.|||||:|||:|||.:.|||||||||||||||.||||||||||||.|.||.|||||||||
Human_SIK2    80 IKLYQVMETKSMLYLVTEYAKNGEIFDYLANHGRLNESEARRKFWQILSAVDYCHGRKIVHRDLKAENLLLDNNMNIKIADFGFGNFFKSGELLATWCGS  179

Fly_SIK2     301 PPYAAPEVFEGKQYTGPEIDIWSLGVVLYVLVCGALPFDGSTLQSLRDRVLSGRFRIPFFMSSECEHLIRRMLVLEPTRRYTIDQIKRHRWMCPEL-LEH  399
                 |||||||||||:||.||::||||:||||||||||||||||.||..||.|||.||||||:|||.:|||||||||||||:|::|.||.|||.|:|||.|: ::.
Human_SIK2   180 PPYAAPEVFEGQQYEGPQLDIWSMGVVLYVLVCGALPFDGPTLPILRQRVLEGRFRIPYFMSEDCEHLIRRMLVLDPSKRLTIAQIKEHKWMLIEVPVQR  279

Fly_SIK2     400 VLIAKYNLGAERQTSV-EPSEDILRIMAEYVGIGSDKTRASLKKNTYDHVAAIYLLLQDRV-SHKKEQSNGLGASALASSTSASRMIYSSRNDHQPTQQQ  497
                 .::  |....|.:.|:|.:|.:||:| ..:||...||..||:..:|:|.||||.||.:|: ||:..                            .|.:|:
Human_SIK2   280 PVL--YPQEQENEPSIGEFNEQVLRLM-HSLGIDQQKTIESLQNKSYNHFAAIYFLLVERLKSHRSS---------------------------FPVEQR  349

Fly_SIK2     498 ---SQQQSKTISTSSILAKDQCHKRLSRHQTVLMSERNAHAGATPTVPDPGPGYYAKYGPLQLPLPLTGHSHLTGYLNGGGVEVDASGIPLPMRYTPLPT  594
                     .|::..||:..::          ::.|||                               .||:|.||
Human_SIK2   350 LDGRQRRPSTIAEQTV----------AKAQTV-------------------------------GLPVTMHS----------------------------  379

Fly_SIK2     545 GPGYYAKYGPLQLPLPLTGHSHLTGYLNGGGVEVDASGIPLPMRYTPLPTAASPAPSNCSSTSSRVGRHSLSSSSPRSHRPVAISLSIDNNPSLANLRCR  644
                              .||:|.||                                                                     |::..||..
Human_SIK2   372 -------------GLPVTMHS---------------------------------------------------------------------PNMRLLRSA  388

Fly_SIK2     645 EMMEAGGGPVGAVGVPLASKQLHQTISEFIIKQSTEDCRALLQQSTAVAEGKDDPPKAESSVGGVPPPASTTPTSSTAGPESGSAPCPGEINGKTIKTMS  744
                 .:.:|..  |.|...|.:..|......|       |:|              .|.||....:....||...         ..|....|
Human_SIK2   389 LLPQASN--VEAFSFPASGCQAEAAFME-------EEC--------------VDTPKVNGCLLDPVPPVLV---------RKGCQSLP-----------  444

Fly_SIK2     745 SSSSFDSKANLGQSFRYKMSAEASKLFQTLQESPLPVEQRTKRRVHVGSTNGSGGDSGQETNDAKSNGDSRSEKKVLAQGSSSTDEGCET--DQGNDPGS  842
                                         |.:.:|                                                     |.|||.||  :...||..
Human_SIK2   445 -----------------------SNMMET-----------------------------------------------------SIDEGLETEGEAEEDPAH  468

Fly_SIK2     843 ASQESKGSNGGGSGNANGGPTSHSSSDLT-RLVGTTTSGQSHKMRSYASSSSSSGVLAGSAGSYSKSLSQNLSRGSSKSNCSGPYDSLDFALPSGKGSLP  941
                 |.:..:        :...|...|:.|::| :||                     |:.|:...:|.:.|.:|....|:.:......|:|...:     |
Human_SIK2   469 AFEAFQ--------STRSGQRRHTLSEVTNQLV---------------------VMPGAGKIFSMNDSPSLDSVDSEYDMGSVQRDLNFLEDN-----P  533

Fly_SIK2     942 SCMGSSSMLAT-PTPASASPAGISSEHSSERSLYGSHNSCIHMPGALPLGLGLPQSSASTPTPNPTPPPNGGGVTFLDKRSPIHFREGRRASDGLVAQGL  1040
                 |.  ...|||. |:|...||. ||..                                  || ||...........:..|||:.||||||||||..:.||:
Human_SIK2   534 SL--KDIMLANQPSPRMTSPF-ISLR----------------------------------PT-NPAMQALSSQKREVHNRSPVSFREGRRASDTSLTQGI  595

Fly_SIK2    1041 LSSGSLLGTSRVYGSYRYEQAKRHGWLEIQQLQQLQQEAAVGHSHPHAHQHPHQHPHPHPQAYGLEELC-QFPNGQ-------FYALPGKHHPLLTLPHH  1132
                 ::          :..:....|:..|.||:.::|.|.|.::  :|         |...|:..|.|..|::|. ..|..:        ...||...||.|:
Human_SIK2   596 VA----------FRQHLQNLARTKGILELNKVQLLYEQ--IG---------PEADPNLAPAAPQLQDLASSCPQEEVSQQQESVSTLPASVHPQLS----  670

Fly_SIK2    1133 HAHPAQH---HHGHHSLFHSGHQATPLLLEAAAGGDMYGHGCYAPPPPPPGLYTHHQLGVGMAVPMSPMQKPPLQQQLLQHRLLQQKRQLFQKQYALEAQ  1229
                    |.|.   .:..|.|     |...||.:|.....:|   |..||                         ..|:|||.:|| |||||...|||..|:|
Human_SIK2   671 ---PRQSLETQYLQHRL-----QKPSLLSKAQNTCQLY---CKEPP-------------------------RSLEQQLQEHR-LQQKRLFLQKQSQLQA-  732

Fly_SIK2    1230 LAGRHHHHQQHSHHHHHHFGHSL-----GPPPPPAPAPEHHLADELYELALLDRPRSGTPRLQHSMTMPGAHAFSQMNLK--------------TSYIS  1309
                     :.:..|.:...:......|     ..|||...||...|...|..:.   .|.|  .::|:|   |....:.|.|:                |....
Human_SIK2   733 ----YFNQMQIAESSYPQPSQQLPLPRQETPPPSQQAPPFSLTQPLSPVL---EPSS--EQMQYS---PFLSQYQEMQLQPLPSTSGPRAAPPLPTQLQQ  820

Fly_SIK2    1310 QSDQVPGAAPNGAGSGGSAAGV-----SCSAP--PSTAPGTPTVKCK-------------PTTPHGHSLDSDYHTSTPVLSLFTPNWQSLVKPLSESPIL  1389
                 |....|...|.....|.:.|.:      :|..|  .|.||..|| .|:             |..|....|.....:.|:                .|||.|.|
Human_SIK2   821 QQPPPPPPPPPRQPGAAPAPLQFSYQTCELPSAASPAPDYPT-PCQYPVDGAQQSDLTGPDCPRSPGLQEAPSSYDPL-------------ALSELPGL  906

Fly_SIK2    1390 EISEHLESV-----------   1398
                 ...|.|::|
Human_SIK2   907 FDCEMLDAVDPQHNGYVLVN    926
```

```
Fly_SIK3      1 MATTPTAGPAAAPPTSSTPQNYKVPSTSKISVDKLLRVGYYELEKTIGKGNFAVVKLATNIVTKTKVAIKIIDKTCLNEEYLNKTFREIAILKSLRHPHI 100
                                                  ...|:| ||::|||||||||||||.||::|||.||| ||||||.:||.|.|.|||:.|:|.|.||||
Human_SIK3    1 -----------------------------------MPARIGYYEIDRTIGKGNFAVVKRATHLVTKAKVAIKIIDKTQLDEENLKKIFREVQIMKMLCHPHI 67

Fly_SIK3    101 TRLYEVMESQSMIYLVTEYAPNGEIFDHLVANGRMKEPEAARVFTQLVSAVHYCHRRGVVHRDLKAENVLLDKDMNIKLADFGFSNHYEEGATLKTWCGS 200
                .|||:|||::.||||||||||..||||||||||:|||.|.||.|.|.|:|:||::||.|.:|||||||||:|||.::||:|||||||.:..|..||||||
Human_SIK3   68 IRLYQVMETERMIYLVTEYASGGEIFDHLVAHGRMAEKEARRKFKQIVTAVYFCHCRNIVHRDLKAENLLLDANLNIKIADFGFSNLFTPGQLLKTWCGS 167

Fly_SIK3    201 PPYAAPEVFQGLEYDGPKSDIWSLGVVLYALVCGALPFDGKTILELKSRVVLGKFRIPFFMSQECEQLIRNMLVVEPDRRYTIKQIIKHRWLSEWQSEMQ 300
                |||||||:|:|.||||||.||||||||||.||||||||||.|:..|::||:.|||||||||||.|||.|||:|||::|:.:::||.||:|:
Human_SIK3  168 PPYAAPELFEGKEYDGPKVDIWSLGVVLYVLVCGALPFDGSTLQNLRARVLSGKFRIPFFMSTECEHLIRHMLVLDPNKRLSMEQICKHKWM-------- 259

Fly_SIK3    301 EQERFGDMSPGSGTVSKSASTSSLGSASDSPPQLDSVVMTHMLQLPGLTADMIAQSVHEQRFDNIYAIYNLLHDKLQQRRRENQRLQHHASLAYSRSRKT 400
                   :.||..|....:  .|....|.......|..:.|::.  ::..||..:...||:....:|:..|||:||.|:  .:|.:..||
Human_SIK3  260 ---KLGDADPNFDRL--IAECQQLKEERQVDPLNEDVLLA--MEDMGLDKEQTLQSLRSDAYDHYSAIYSLLCDR--HKRHKTLRL-------------- 336

Fly_SIK3    401 SITTGVVDRSEPVKQESLDRLSPLSNANASSSALGFGWSDVAVDL--------EKYGDFELEC-LARSNDPPVNAQHLSAHAGGGVNGANTRRHTVGPGD 491
                    |.:    |....:|...:|: |..|..:.......|...|.|        |..|...|:. ......|....::||           .||||||..|
Human_SIK3  337 ----GAL----PSMPRALAFQAPV-NIQAEQAGTAMNISVPQVQLINPENQIVEPDGTLNLDSDEGEEPSPEALVRYLS-----------MRRHTVGVAD 416

Fly_SIK3    492 VAHE-----QALANPNVPPIDFKCPPQCSDPTQPVPYYPVNLPMLQNQPLHNLTIKDQHLLKPPVVM---GASSFGRRASDGGANLHIYYPATGTVVGPA 583
                ...|     |.|. |..|.:.:.|.....|.  |.:.....|||...||...|..|:|.||:||.:.   |....|||||||||:.::
Human_SIK3  417 PRTEVMEDLQKLL-PGFPGVNPQAPFLQVAPN--VNFMHNLLPMQNLQPTGQLEYKEQSLLQPPTLQLLNGMGPLGRRASDGGANIQLH----------- 502

Fly_SIK3    584 QGQQMDTAGYYINPNCGTDPL-----AVQELSPLNEQSVAQMQCCQENATGECNEE-LQSYMQKRGGTQRHTVGCTEDLSVAHGSGAGSQS--------Q 669
                 .||:      :....|..||     ||..::|::|            |:.||.::| :||...|...|....   ||..|.......|:.|         :
Human_SIK3  503 -AQQL------LKRPRGPSPLVTMTPAVPAVTPVDE----------ESSDGEPDQEAVQSSTYKDSNTLHLP---TERFSPVRRFSDGAASIQAFKAHLE 582

Fly_SIK3    670 APTTSSNMRQRRTGLLTVTERPPGALERLTKIL-------------------------------------------------------------------- 702
                ....:|:::|.:.             ..|:|.|:.
Human_SIK3  583 KMGNNSSIKQLQQ-----------ECEQLQKMYGGQIDERTLEKTQQQHMLYQQEQHHQILQQQIQDSICPPQPSPPLQAACENQPALLTHQLQRLRIQP 671

Fly_SIK3    703 ---------------------------------------------------------------------------------------------------- 702

Human_SIK3  672 SSPPPNHPNNHLFRQPSNSPPPMSSAMIQPHGAASSSQFQGLPSRSAIFQQQPENCSSPPNVALTCLGMQQPAQSQQVTIQVQEPVDMLSNMPGTAAGSS 771

Fly_SIK3    703 ---------------------------------------------------------------------------------------------------- 702

Human_SIK3  772 GRGISISPSAGQMQMQHRTNLMATLSYGHRPLSKQLSADSAEAHSLNVNRFSPANYDQAHLHPHLFSDQSRGSPSSYSPSTGVGFSPTQALKVPPLDQFP 871

Fly_SIK3    703 ---------------------------------------------------------------------------------------------------- 702

Human_SIK3  872 TFPPSAHQQPPHYTTSALQQALLSPTPPDYTRHQQVPHILQGLLSPRHSLTGHSDIRLPPTEFAQLIKRQQQQRQQQQQQQQQQEYQELFRHMNQGDAGS 971

Fly_SIK3    703 ---------------------------------------------------------------------------------------------------- 702

Human_SIK3  972 LAPSLGGQSMTERQALSYQNADSYHHHTSPQHLLQIRAQECVSQASSPTPPHGYAHQPALMHSESMEEDCSCEGAKDGFQDSKSSSTLTKGCHDSPLLLS 1071

Fly_SIK3    703 ---------------------------------------------------------------------------------------------------- 702

Human_SIK3 1072 TGGPGDPESLLGTVSHAQELGIHPYGHQPTAAFSKNKVPSREPVIGNCMDRSSPGQAVELPDHNGLGYPARPSVHEHHRPRALQRHHTIQNSDDAYVQLD 1171

Fly_SIK3    703 -------------------------------------------------------------------------------------------    702

Human_SIK3 1172 NLPGMSLVAGKALSSARMSDAVLSQSSLMGSQQFQDGENEECGASLGGHEHPDLSDGSQHLNSSCYPSTCITDILLSYKHPEVSFSMEQAGV    1263
```
