## Supplementary Table 1 for "Salt Inducible Kinases as Novel Notch Interactors in the Developing *Drosophila* Retina"

| Genotype | Sensitized background (*ey-Gal4* > *Dl*) | | | | | | Eyeful background (*ey-Gal4* > *Dl, GS88A8*) | | | | | |
|---|---|---|---|---|---|---|---|---|---|---|---|---|
| | Not affected / Normal | Folding / Overgrowth | Ectopic eyes | Eye loss | Metastasis | Total affected | Not affected / Normal | Folding / Overgrowth | Ectopic eyes | Eye loss | Metastasis | Total affected |
| - | 98.6 | 1.1 | 0.3 | 0.0 | 0.0 | **1.4 ±0.3** | 71.4 | 19.8 | 5.7 | 1.8 | 1.4 | **28.6 ±11.0** |
| >*Sik2 OE* | 76.7 | 19.1 | 4.2 | 0.0 | 0.0 | **23.3 ±3.5** | 67.8 | 13.9 | 10.0 | 6.2 | 2.1 | **32.1 ±10.3** |
| >*Sik2*$^{RNAi}$ | 87.9 | 4.3 | 5.1 | 2.5 | 0.2 | **12.1 ±3.2** | 52.7 | 29.5 | 8.4 | 8.1 | 1.3 | **47.3 ±10.1** |
| >*Sik2*$^{S1032A}$ | 92.3 | 7.1 | 0.6 | 0.0 | 0.0 | **7.7 ±0.5** | 76.4 | 8.8 | 7.2 | 4.4 | 3.2 | **23.6 ±4.6** |
| >*Sik2*$^{K170M}$ | 78.1 | 19.4 | 2.0 | 0.6 | 0.0 | **21.9 ±3.0** | 67.1 | 17.6 | 45.0 | 6.6 | 3.9 | **33.0 ±10.1** |
| >*Sik3 OE* | 97.9 | 1.5 | 0.4 | 0.0 | 0.2 | **2.1 ±0.9** | 74.5 | 15.0 | 4.8 | 4.9 | 1.0 | **25.5 ±9.2** |
| >*Sik3*$^{RNAi}$ | 92.4 | 7.6 | 0.0 | 0.0 | 0.0 | **7.6 ±1.0** | 61.5 | 23.9 | 11.9 | 2.6 | 0.2 | **38.5 ±11.9** |
| >*Sik3*$^{S563A}$ | 60.5 | 9.0 | 12.6 | 17.0 | 0.9 | **39.5 ±6.1** | 55.0 | 21.0 | 9.0 | 14.0 | 1.0 | **45.0 ±6.1** |

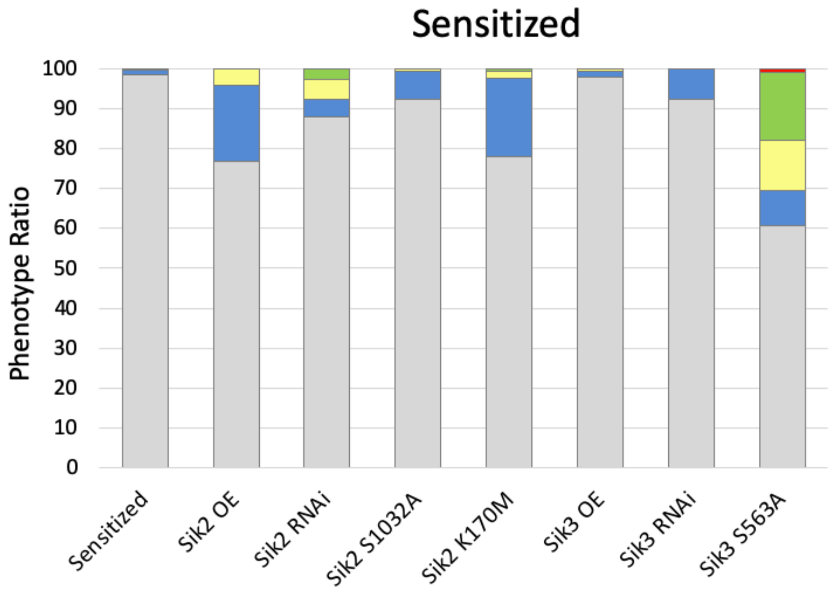


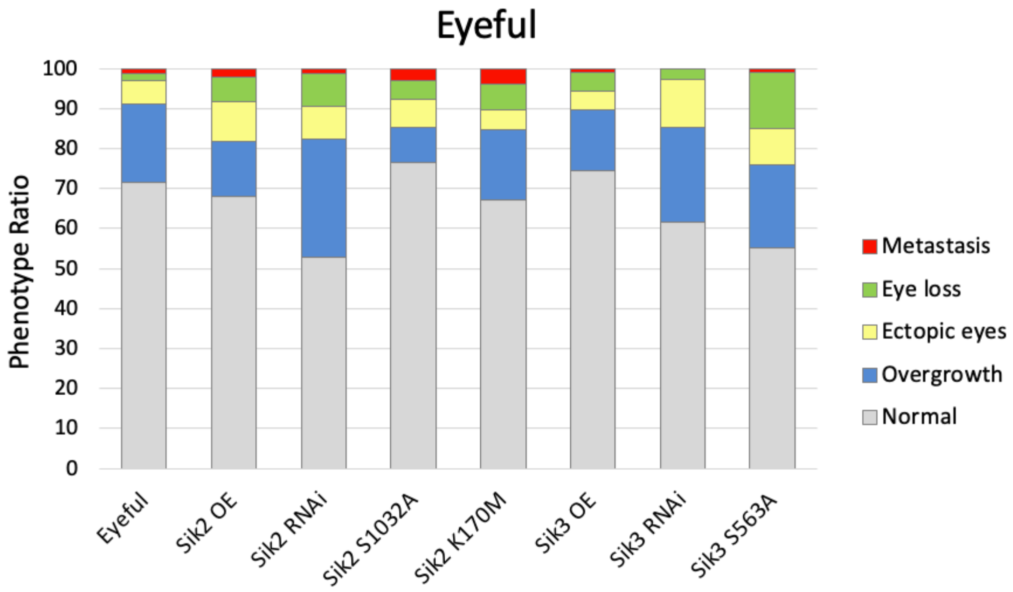
