## Supplementary Table 2 for "Salt Inducible Kinases as Novel Notch Interactors in the Developing *Drosophila* Retina"

| Below are the full genotypes belonging the figures mentioned: | | | |
| --- | --- | --- | --- |
| **Figure** | **Frame** | **Name** | **Full genotype** |
| **Figure 1** | Fig. 1H - sensitized | - | *ey-GAL4 , UAS Delta / +* |
|  |  | *>Sik2 OE* | *ey-GAL4 , UAS Delta / + ; UAS-Sik2 / +* |
|  |  | *>Sik2^RNAi^* | *ey-GAL4 , UAS Delta / UAS-Sik2^RNAi^* |
|  |  | *>Sik2^S1032A^* | *ey-GAL4 , UAS Delta / + ; UAS-Sik2^S1032A^/ +* |
|  |  | *>Sik2^K170M^* | *ey-GAL4 , UAS Delta / UAS-Sik2^K170M^* |
|  |  | *>Sik3 OE* | *ey-GAL4 , UAS Delta / + ; UAS-Sik3::T2A::mCherry / +* |
|  |  | *>Sik3^RNAi^* | *ey-GAL4 , UAS Delta / + ; UAS-Sik3^RNAi^ / +* |
|  |  | *>Sik3^S563A^* | *ey-GAL4 , UAS Delta / UAS-Sik3^S563A^* |
|  | Fig. 1H - eyeful | - | *ey-GAL4 , UAS Delta, GS88A8 / +* |
|  |  | *>Sik2 OE* | *ey-GAL4 , UAS Delta, GS88A8 / + ; UAS-Sik2 / +* |
|  |  | *>Sik2^RNAi^* | *ey-GAL4 , UAS Delta, GS88A8 / UAS-Sik2^RNAi^* |
|  |  | *>Sik2^S1032A^* | *ey-GAL4 , UAS Delta, GS88A8 / + ; UAS-Sik2^S1032A^/ +* |
|  |  | *>Sik2^K170M^* | *ey-GAL4 , UAS Delta, GS88A8 / UAS-Sik2^K170M^* |
|  |  | *>Sik3 OE* | *ey-GAL4 , UAS Delta, GS88A8 / + ; UAS-Sik3::T2A::mCherry / +* |
|  |  | *>Sik3^RNAi^* | *ey-GAL4 , UAS Delta, GS88A8 / + ; UAS-Sik3^RNAi^ / +* |
|  |  | *>Sik3^S563A^* | *ey-GAL4 , UAS Delta, GS88A8 / UAS-Sik3^S563A^* |
| **Figure 2** | Fig. 2A-A′′ |  | *UAS-Dicer2 / + ; ey-GAL4, lGMR-GAL4 / +* |
|  | Fig. 2B-B′′ | *>Sik2 OE* | *UAS-Dicer2 / + ; ey-GAL4, lGMR-Gal4 / + ; UAS-Sik2 / +* |
|  | Fig. 2C-C′′ | *>Sik2^RNAi^* | *UAS-Dicer2 / + ; ey-GAL4, lGMR-GAL4 / UAS-Sik2^RNAi^* |
|  | Fig. 2D | *ey>Sik2^RNAi^* | *ey-GAL4 / + ; UAS-Sik2^RNAi^* |
|  | Fig. 2E-E′′ | *>Sik3 OE* | *UAS-Dicer2 / + ; ey-GAL4, lGMR-GAL4 / + ; UAS-Sik3::T2A::mCherry / +* |
|  | Fig. 2F-F′′ | *>Sik3^RNAi^* | *UAS-Dicer2 / + ; ey-GAL4, lGMR-GAL4 / + ; UAS-Sik3^RNAi^ / +* |
|  | Fig. 2G-G′′ | *>Sik2^K170M^* | *UAS-Dicer2 / + ; ey-GAL4, lGMR-GAL4 / + ; UAS-Sik2^K170M^ / +* |
|  | Fig. 2H-H′′ | *>Sik2^S1032A^* | *UAS-Dicer2 / + ; ey-GAL4, lGMR-GAL4 / + ; ; UAS-Sik2^S1032A^ / +* |
| **Figure 3** | Fig. 3A | - | *UAS-Dicer2 / + ; ey-GAL4, lGMR-GAL4 , UAS-Fng/ +* |
|  | Fig. 3B | *>Sik2^K170M^* | *UAS-Dicer2 / + ; ey-GAL4, lGMR-GAL4 , UAS-Fng/ UAS-Sik2^K170M^* |
|  | Fig. 3C | *>Sik3^S563A^* | *UAS-Dicer2 / + ; ey-GAL4, lGMR-GAL4 , UAS-Fng/ UAS-Sik3^S563A^* |
|  | Fig. 3D | - | *UAS-Dicer2 / + ; ey-GAL4, lGMR-GAL4 / + ; UAS-Ser^RNAi^ / +* |
|  | Fig. 3E | *>Sik2 OE* | *UAS-Dicer2 / + ; ey-GAL4, lGMR-GAL4 / + ; UAS-Ser^RNAi^ / + ; UAS-Sik2 / +* |
|  | Fig. 3F | *>Sik2^K170M^* | *UAS-Dicer2 / + ; ey-GAL4, lGMR-GAL4 / + ; UAS-Ser^RNAi^ / UAS-Sik2^K170M^* |
|  | Fig. 3G | *>Sik2^S1032A^* | *UAS-Dicer2 / + ; ey-Gal4, lGMR-GAL4 / + ; UAS-Ser^RNAi^ / + ; UAS-Sik2^S1032A^/ +* |
|  | Fig. 3H | *>Sik3 OE* | *UAS-Dicer2 / + ; ey-GAL4, lGMR-GAL4 / + ; UAS-Ser^RNAi^ /+ ; UAS-Sik3::T2A::mCherry / +* |
|  | Fig. 3I | *>Sik3^S563A^* | *UAS-Dicer2 / + ; ey-GAL4, lGMR-GAL4 / + ; UAS-Ser^RNAi^ /UAS-Sik3^S563A^* |
|  | Fig. 3J | - | *UAS-Dicer2 / UAS-Delta ; ey-GAL4, lGMR-GAL4 / +* |
|  | Fig. 3K | *>Sik2^S1032A^* | *UAS-Dicer2 / UAS-Delta ; ey-GAL4, lGMR-GAL4 / + ; UAS-Sik2^S1032A^/ +* |
|  | Fig. 3L | *>Sik2^K170M^* | *UAS-Dicer2 / UAS-Delta ; ey-GAL4, lGMR-GAL4 / UAS-Sik2^K170M^* |
|  | Fig. 3M | *>Sik3^RNAi^* | *UAS-Dicer2 / UAS-Delta ; ey-GAL4, lGMR-GAL4 / UAS-Sik3^RNAi^* |
|  | Fig. 3N | *>Sik3^S563A^* | *UAS-Dicer2 / UAS-Delta ; ey-GAL4, lGMR-GAL4 / UAS-Sik3^S563A^* |
|  | Fig. 3O | *>Sik2^RNAi^* | *UAS-Dicer2 / UAS-Delta ; ey-GAL4, lGMR-GAL4 /+ ; UAS-Sik2^RNAi^ / +* |
| **Supp. Figure 2** | S. Fig. 2C | Control | *w^1118^* |
|  |  | Sik3::T2A::mCherry | *ey-GAL4, lGMR-GAL4 / + ; UAS-Sik3::T2A::mCherry / +* |
| **Supp. Figure 3** | S. Fig. 3A | Control-hid | *neoFRT42D, GMR-hid* / *+ ; ey-Gal4, UAS-Flp* / *+.* |
|  | S. Fig. 3A′ | Control-Flp | *CyO* / *+ ; ey-Gal4, UAS-Flp* / *+* |
|  | S. Fig. 3A′′ | *Sik3^∆109^* | Eye: *neoFRT42D Sik3^∆109^ ; ey-Gal4, UAS-Flp/+*  Rest of body: *neoFRT42D Sik3^∆109^ / neoFRT42D, GMR-hid ; ey-Gal4, UAS-Flp/+* |
|  | S. Fig. 3B | MARCM Control | *ey-Flp / + ; neoFRT42D, GMR-myrGFP / +* |
|  | S. Fig. 3B′ | *Sik3^∆109^* MARCM | Red regions: *ey-Flp / + ; neoFRT42D, Sik3^∆109^* / *neoFRT42D, GMR-myrGFP*  White regions: *ey-Flp / + ; neoFRT42D, Sik3^∆109^* |
| **Supp. Figure 4** | S. Fig. 5A | Control | *actin-GAL4 / CyO* |
|  | S. Fig. 5B | *>Sik2^K170M^* | *actin-GAL4 / UAS-Sik2^K170M^* |
|  | S. Fig. 5C | *>Sik2^S1032A^* | *actin-GAL4 / UAS-Sik2^S1032A^* |
|  | S. Fig. 5D | *>Sik2 OE* | *mirr-GAL4 / UAS-Sik2* |
|  | S. Fig. 5E | *>Sik3^S563A^* | *UAS-Sik3^S563A^ / + ; mirr-GAL4 / +* |
|  | S. Fig. 5F | *>Delta^DN^* | *sca-Gal4 / + ; UAS-Delta^DN^ / +* |
